# A biological light sheet links cellular voltage to mesoscopic brain dynamics

**DOI:** 10.64898/2026.10.30.761751

**Authors:** Tsai-Wen Chen, Bei-Jung Lin

## Abstract

Membrane voltage is the fundamental currency of neural computation, yet cellular-resolution voltage imaging of large neuronal populations *in vivo* remains limited by trade-offs among imaging speed, field of view, and optical sectioning. We overcome this challenge by encoding a biological light sheet in the brain: Selective Plane Expression (SPress) restricts fluorescent indicator expression to a user-defined, 20-μm-thick plane, suppressing background at its source. Following one-time two-photon photoactivation, SPress enables standard widefield microscopes to capture high-contrast kilohertz voltage dynamics from hundreds of densely labeled neurons across millimeter-scale cortical sheets and columns *in vivo*. These recordings reveal mesoscopic subthreshold waves that flexibly reorganize population-wide spike timing, together with cellular-scale wavefront microstructure that preserves stereotyped local recruitment biases. By shifting optical sectioning from instrumentation to gene expression, SPress makes kilohertz cellular-resolution voltage imaging accessible across mesoscopic scales.

## Main Text

Brain function emerges from large-scale patterns of neural activity. Mesoscopic brain dynamics—including cortical states, traveling waves, and oscillatory rhythms spanning hundreds of micrometers to millimeters (*1–3*)—are central to understanding behavior, computation, and disease. These patterns must ultimately arise from the membrane-potential dynamics of individual neurons, yet how cellular voltage is coordinated across mesoscopic scales, and how this organization shapes spatiotemporal patterns of neuronal spiking, remain largely unknown.

Voltage imaging offers a promising route to addressing these questions (*4–6*). By converting membrane voltage into changes in fluorescence, voltage indicators can report activity from large neuronal populations with millisecond precision. In the living brain, however, neurons are embedded in complex three-dimensional tissue, making it difficult to isolate cellular voltage signals from surrounding out-of-focus fluorescence.

Conventional approaches achieve optical sectioning by restricting fluorescence excitation or detection to a thin focal plane, as in confocal (*7*) and multiphoton microscopy (*8*). Although these scanning-based methods have traditionally been limited in speed, recent innovations have extended optical sectioning into the kilohertz regime for voltage imaging (*9–13*). These techniques represent substantial engineering advances but require complex instrumentation and remain constrained by a trade-off between imaging speed and field of view (*14*) (fig. S1). Consequently, kilohertz cellular voltage imaging with millimeter-scale coverage remains difficult for existing optical-sectioning approaches (*14*) (fig. S1). By contrast, single-photon widefield microscopy offers simplicity, speed, and spatial coverage but lacks optical sectioning, making cellular voltage signals difficult to resolve in densely labeled tissue (*15, 16*).

We therefore asked whether optical sectioning must be provided by the microscope at all. Instead of suppressing out-of-focus fluorescence optically, could background be minimized at its source by restricting indicator expression to a thin, user-defined plane? Such an approach would encode optical sectioning in the sample itself, bypassing this scanning-imposed trade-off while preserving the speed, field of view, and accessibility of widefield microscopy.

Previous strategies for layer-restricted labeling—including in utero electroporation (*17*), Cre-driver lines (*18*), and enhancer-based viral targeting (*19*)—target developmental, genetic, or cell-type identities rather than a geometrically defined optical plane. Consequently, labeled regions are often hundreds of micrometers thick rather than confined to an optical-section-like plane. Moreover, expression cannot be flexibly positioned at user-defined depths or orientations to match the geometry of the target structure under study.

### Selective Plane Expression creates a biological light sheet

To address this challenge, we developed SPress, a genetic strategy that restricts expression to a user-defined, ∼20-µm-thick plane. SPress uses a photoactivatable recombination system (*20*), activated by scanning the focal volume of a conventional two-photon microscope across the target plane (Fig. 1A). This one-time expression-patterning step selectively induces recombination within the illuminated layer, stably activating fluorescent reporter expression in a thin sheet of cells.

**Fig. 1.**
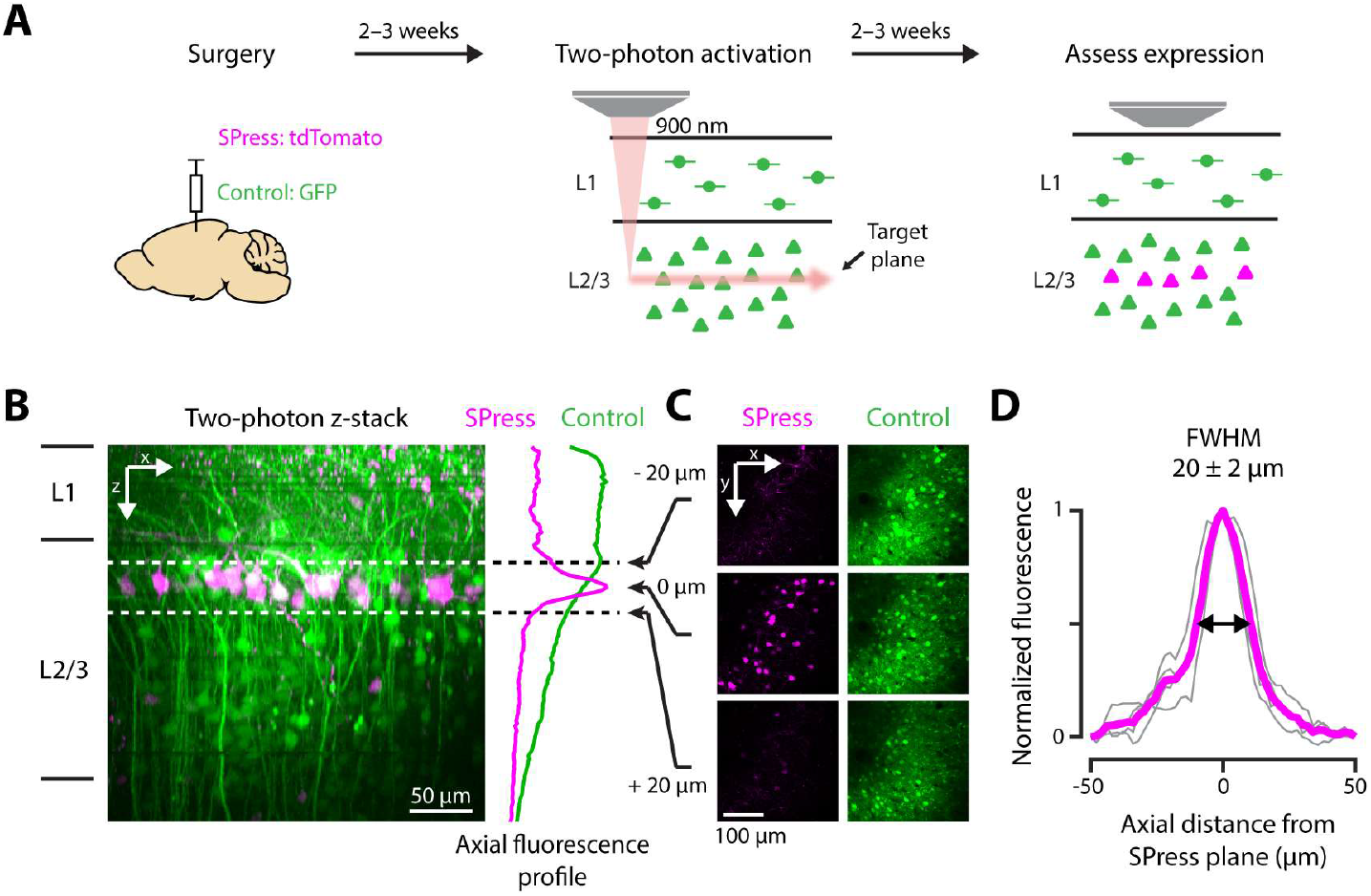
Selective Plane Expression creates a biological light sheet. (**A**) SPress labeling strategy. AAVs carrying a photoactivatable Cre recombinase (iCreV), Cre-dependent tdTomato (SPress; magenta), and constitutive GFP (green) were injected during cranial window surgery (left), followed 2–3 weeks later by two-photon activation at 900 nm in a defined cortical plane (middle). Expression was assessed 2–3 weeks later by two-photon z-stack imaging (right). (**B**) Side-view two-photon image of cortical fluorescence after SPress induction, together with axial fluorescence profiles for SPress (magenta) and control GFP (green). Dashed lines mark the targeted SPress plane. (**C**) Images acquired at three axial positions relative to the targeted plane (−20, 0, and +20 µm), shown for SPress (magenta) and control GFP (green). (**D**) Normalized axial fluorescence profile for SPress, with full width at half maximum (FWHM) indicated and reported as mean ± SD. Gray lines show individual animals (n = 4), and magenta shows the mean.

We tested SPress in the mouse cerebral cortex by injecting adeno-associated viruses encoding iCreV (*20*), a photoactivatable Cre recombinase, together with a Cre-dependent tdTomato reporter during cranial window surgery. An additional AAV constitutively expressing GFP labeled the injection site and served as a non-SPress control. After viral expression, two-photon scanning at 900 nm was used to activate recombination within a user-defined optical plane located ∼100–200 µm below the cortical surface, targeting layer 2/3 pyramidal neurons. Two weeks later, SPress-mediated tdTomato expression and GFP control expression were examined by two-photon microscopy.

SPress restricted tdTomato expression to a thin cortical sheet centered on the activation plane. In an example x–z section of an SPress-labeled region (Fig. 1B), tdTomato fluorescence was tightly confined to the activated layer and fell sharply above and below it (Fig. 1C). Quantification of mean fluorescence intensity (Fig. 1D) and labeled-cell number (fig. S2) as a function of depth revealed a narrow peak, with a full width at half maximum of 20 ± 2 µm (mean ± SD; n = 4 animals), consistent with confinement set primarily by the spatial restriction of two-photon activation. In contrast, control GFP expression was broadly distributed, producing a substantially wider fluorescence profile across depth (Fig. 1B, green). These results show that SPress confines fluorescence to a thin plane, effectively creating a biological light sheet within the brain.

### Optical-sectioning-like voltage imaging with widefield microscopy

Voltage imaging is highly sensitive to out-of-plane background because it relies on detecting small, fast fluorescence changes from individual neurons. The tight spatial confinement of SPress should minimize this background, enabling optical-sectioning-like contrast. To test this idea, we combined SPress with the chemigenetic voltage indicator Voltron2 (*21*) and targeted expression to a defined plane in layer 2/3 of mouse cortex (Fig. 2A). A constitutively expressed GFP reporter in the same mouse and field of view served as a non-SPress control. Two weeks after SPress activation, we imaged Voltron2 and GFP fluorescence by widefield microscopy and acquired two-photon z-stacks of both signals to quantify their axial expression profiles.

**Fig. 2.**
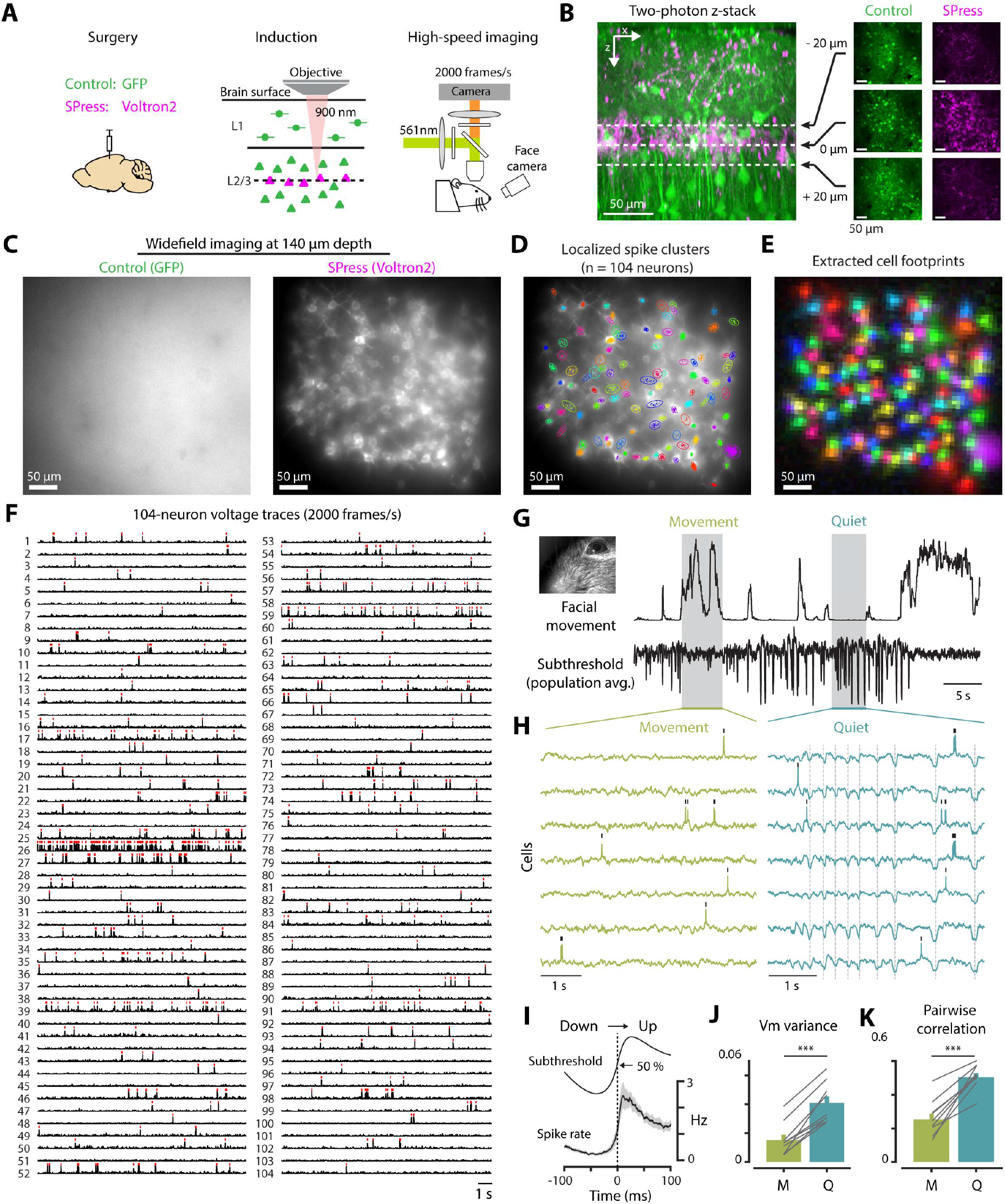
Optical-sectioning-like voltage imaging with widefield microscopy. (**A**) Experimental configuration. AAVs encoding SPress (Voltron2; magenta) and control (GFP; green) were injected (left), followed 2-3 weeks later by two-photon induction (900 nm) at a defined cortical depth. Widefield voltage imaging was performed another 2-3 weeks later at 2000 frames per second. **(B**) Side-view two-photon z-stack showing tight axial confinement of SPress (magenta) relative to control expression (green). Right, images acquired at three axial positions (−20, 0, and +20 µm) relative to the targeted plane. (**C**) Widefield images (∼140 µm depth) comparing control GFP fluorescence (left) and SPress Voltron2 labeling (right) in the same field of view. (**D**) Detected spikes overlaid on the widefield image. Colored outlines indicate distinct spike clusters (104 clusters). (**E**) Extracted cells displayed as color-coded spatial footprints. (**F**) Extracted voltage traces from 104 simultaneously imaged cells at 2000 frames per second, high-pass filtered to highlight spiking activity. Red ticks indicate detected spikes. (**G**) Facial movement (top; extracted from video recordings of the animal’s face, inset) and population-averaged subthreshold voltage signal (bottom). Shaded regions indicate example movement and quiet periods. (**H**) Example single-cell voltage traces during movement (left) and quiet (right) periods. **(I)** Subthreshold voltage signal (top) and spike rate (bottom) aligned to hyperpolarized-to-depolarized transitions and averaged across neurons (n = 1,686 cells from 9 animals) and events (n = 879). Gray shaded area indicates s.e.m. (**J**) Variance of subthreshold signal during movement (M) and quiet (Q) periods. (**K**) Pairwise correlation of subthreshold signal during movement (M) and quiet (Q) periods. Each point in (J) and (K) shows the mean value per animal; lines connect paired measurements (n = 9 mice, ***P < 0.001, two-sided paired t-test).

GFP-labeled cells were broadly distributed across depth (Fig. 2B, green), producing diffuse fluorescence with no discernible cell bodies in widefield images (Fig. 2C, left). In contrast, SPress restricted Voltron2 expression to a thin fluorescent sheet (Fig. 2B, magenta, Movies S1 and S2), allowing densely labeled neuronal somata located >100 µm below the cortical surface to be readily resolved with a standard widefield microscope (Fig. 2C, right).

The high contrast provided by SPress enabled high-quality, kilohertz voltage imaging from the labeled neuronal sheet. Within a ∼300 × 300 µm field of view, voltage signals from >100 neurons were recorded simultaneously at 2 kHz (Fig. 2D-F). To minimize the effects of emission scattering at depths beyond ∼100 µm, we combined SPress with Activity Localization Imaging (ALI) (*16*), a spike-localization-based method for separating signals from neighboring neurons (fig. S3). Together, SPress and ALI allowed high-SNR detection of neuronal spikes (Fig. 2F) and subthreshold voltage fluctuations (fig. S4) from depths up to ∼200 µm below the cortical surface (fig. S5; SNR = 7.8 ± 1.0; photobleaching = 5.1 ± 1.1% min^−1^; mean ± s.d.; n = 1,686 cells from 9 animals).

The extracted optical voltage signals exhibited characteristic features of cortical membrane potential dynamics. During quiet wakefulness, neurons showed slow, large-amplitude, synchronized depolarizing and hyperpolarizing transitions (Fig. 2G and H, “Quiet”) consistent with cortical up and down states (*22, 23*). Spiking was sparse and occurred predominantly during depolarized states, while being largely suppressed during hyperpolarized periods (Fig. 2H and I). These slow subthreshold fluctuations were strongly modulated by ongoing behavior, being prominent during quiet wakefulness and markedly reduced during facial movement (Fig. 2G and H, “Movement”). Both the amplitude of subthreshold voltage fluctuations and the correlation between cells were markedly reduced during movement (Fig. 2J and K), consistent with known behavior-dependent changes in cortical membrane potential dynamics (*23, 24*).

### Mesoscopic subthreshold waves flexibly organize spike timing

How membrane voltage dynamics of individual neurons are coordinated across large cortical areas remains poorly understood. Across neurons, cellular voltage could fluctuate synchronously (*24, 25*), follow stereotyped recruitment sequences (*26, 27*), or exhibit event-dependent organization across cortical space (*28, 29*). Testing these possibilities requires simultaneous measurements of subthreshold voltage and spiking from individual neurons across millimeter-scale cortical areas, a capability not readily achieved with existing methods.

Because SPress achieves optical-sectioning-like contrast through selective expression, the imaging area could be scaled by extending the labeled plane. To demonstrate this capability, we injected AAVs at multiple sites across the cranial window and activated SPress over a 1 × 1 mm cortical sheet targeting pyramidal neurons (Fig. 3A, see Methods). This produced a millimeter-scale sheet of Voltron2-labeled neurons that could be imaged *in vivo* at kilohertz speed with high contrast across the entire field using a standard widefield microscope (Fig. 3B–D). Individual neurons were readily visible throughout the field of view (Fig. 3C, Movie S3), and extracted spike locations aligned accurately with neuronal positions in widefield images (Fig. 3C and D).

**Fig. 3.**
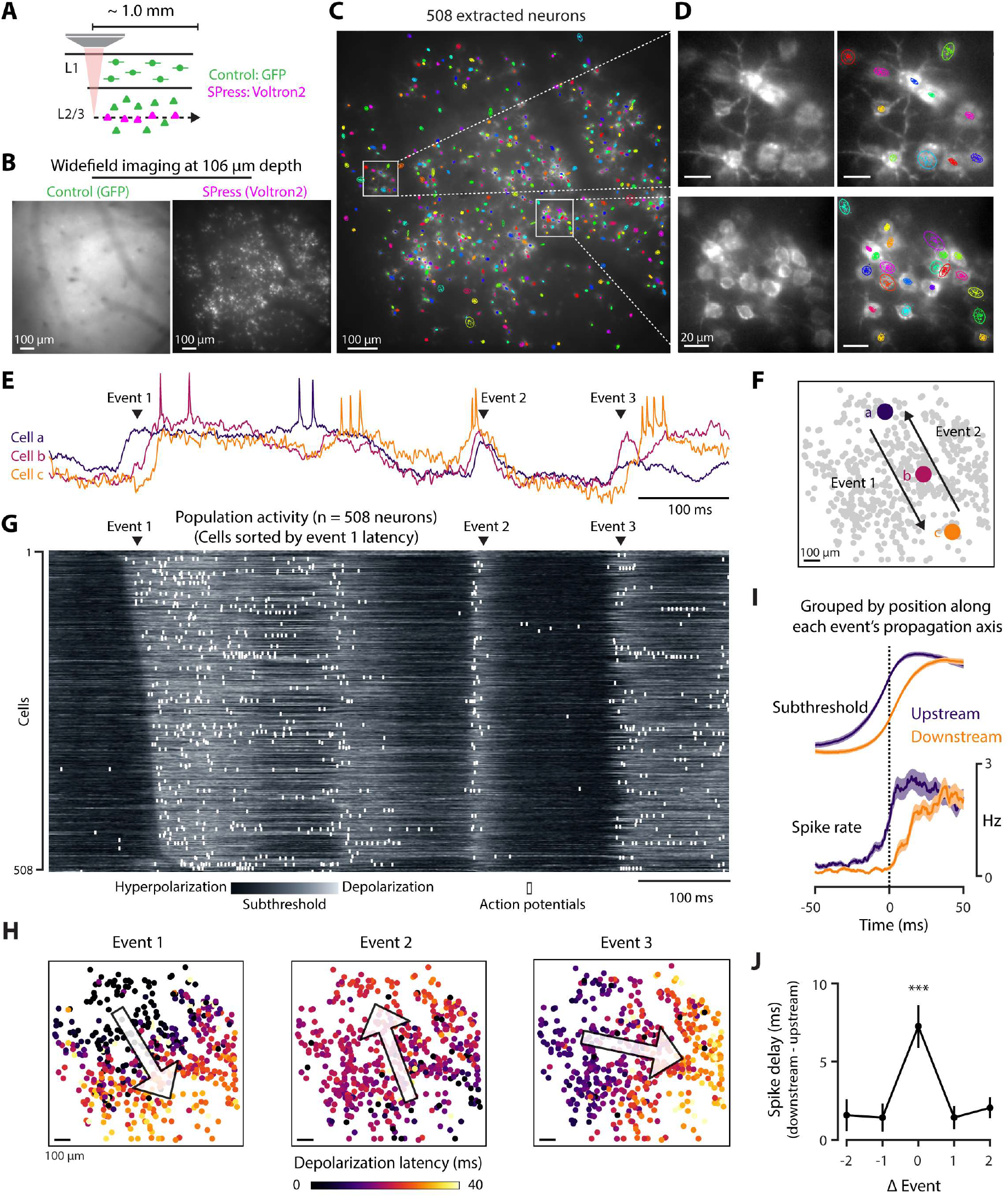
Mesoscopic subthreshold waves flexibly organize spike timing. **(A)** AAVs encoding control GFP and SPress Voltron2 were delivered across a broad cortical region, and two-photon induction was performed along a cortical plane spanning ∼1.0 × 1.0 mm, targeting layer 2/3 pyramidal neurons. **(B)** Widefield images of control GFP (left) and SPress Voltron2 (right) acquired at 106 µm depth from the same cortical field of view. **(C)** Extracted cells (n = 508) overlaid on the widefield image. Boxed regions are shown at higher magnification in (D). **(D)** Enlarged views showing widefield fluorescence (left) and corresponding spike clusters (right) in local regions of the field of view. Colored contours indicate distinct spike clusters. **(E)** Example voltage traces from three neurons (cells a–c, indicated in F). Arrowheads mark three example depolarizing events. **(F)** Spatial positions of neurons a–c within the imaged population. Arrows illustrate the propagation direction of two example events. **(G)** Population activity from all 508 neurons. Rows correspond to individual neurons, sorted by depolarization latency in Event 1. Grayscale indicates subthreshold membrane potential, and white ticks indicate action potentials. **(H)** Spatial maps of depolarization latency for the three example events shown in (E) and (G). Each dot represents one neuron, colored by its depolarization latency within the event. Arrows indicate the fitted propagation direction for each event. **(I)** Subthreshold membrane potential (top) and spike rate (bottom) of neurons grouped by spatial position along the event-specific propagation axis. Upstream (purple) and downstream (orange) cells were defined independently for each event as neurons located on the earlier- and later-arriving sides of the fitted wavefront, respectively. Signals were aligned to the Down-to-Up state transition and averaged across cells and events. Shaded areas indicate s.e.m. **(J)** Spike timing delay quantified by applying the upstream/downstream grouping from each event to the same event or to nearby events. Delay was defined as the difference in the time to half-maximal spike-rate increase between downstream and upstream cells. Error bars indicate s.e.m. (n = 9 animals). ***P < 0.001, two-sided t-test comparing Δ = 0 with the mean of the nonzero offsets (Δ = −2, −1, +1, +2).

Simultaneous voltage imaging from >500 neurons at 2000 frames per second revealed large-scale coordination of cortical membrane-potential dynamics. Large depolarizing events were often shared across widely separated neurons (Fig. 3E–G), indicating coordinated activity across the population.

These events, however, were not globally synchronous. Within each event, neurons depolarized at different times, and the recruitment sequence varied from event to event (Fig. 3E). In some cases, the same neurons were recruited in opposite temporal orders during different events (Fig. 3E–G, events 1 and 2), showing that cellular voltage dynamics do not follow a fixed activation sequence but instead propagate through flexible, event-specific trajectories. Latency maps across neurons revealed two-dimensional propagation of subthreshold depolarization across the cortical sheet, with different events spreading in distinct directions (Fig. 3H).

Previous studies have identified reproducible firing sequences across neuronal populations (*27, 30, 31*). We asked whether event-to-event changes in subthreshold-wave direction could flexibly reorder firing among the same neurons. For each depolarizing event, neurons were classified as upstream or downstream according to their position along the fitted propagation axis. As expected, upstream neurons depolarized earlier and, importantly, showed an earlier rise in spike rate than downstream neurons (Fig. 3I), indicating that wave propagation across the cortical sheet shaped spiking output.

We then asked whether this spike-timing difference was tied to each event’s propagation axis or reflected fixed early-and late-spiking populations. Across nine animals, the delay was greatest for the defining event and markedly reduced when the same upstream/downstream grouping was applied to neighboring events (P < 0.001; Fig. 3J). Thus, firing order was not fixed but dynamically assembled by each subthreshold wave’s propagation trajectory. Together, these results show that millimeter-scale subthreshold waves flexibly reorganize neuronal firing within the same cortical network.

### Vertical SPress reveals event-specific recruitment of cortical layers

The cortex is organized into columns and layers with distinct cell types, connectivity, and activity patterns (*32, 33*). Large-scale cortical waves are often measured from the brain surface (*2*), but how these mesoscopic dynamics engage the layered architecture of cortex remains poorly understood. In particular, surface measurements alone cannot determine whether cortical recruitment exhibits a stereotyped deep-to-superficial bias (*34*) or follows event-specific trajectories spanning both cortical depth and tangential position.

To access this geometry, we used an implanted microprism (*35*) to image voltage dynamics across cortical depth. Implanted optical interfaces such as microprisms (*35*) and GRIN lens microendoscopes (*36*) are widely used to access brain regions or viewing geometries that are otherwise difficult to image. However, direct widefield imaging through these preparations is often compromised by out-of-focus background. The flexibility of SPress made it possible to orient the expression plane vertically (Fig. 4A), creating a thin sheet of Voltron2-labeled neurons parallel to the prism surface that could be imaged with high contrast using a standard widefield microscope (Fig. 4B). This configuration enabled simultaneous voltage imaging from large populations of layer 2/3 and layer 5 neurons without specialized optical-sectioning hardware (Fig. 4B).

**Fig. 4.**
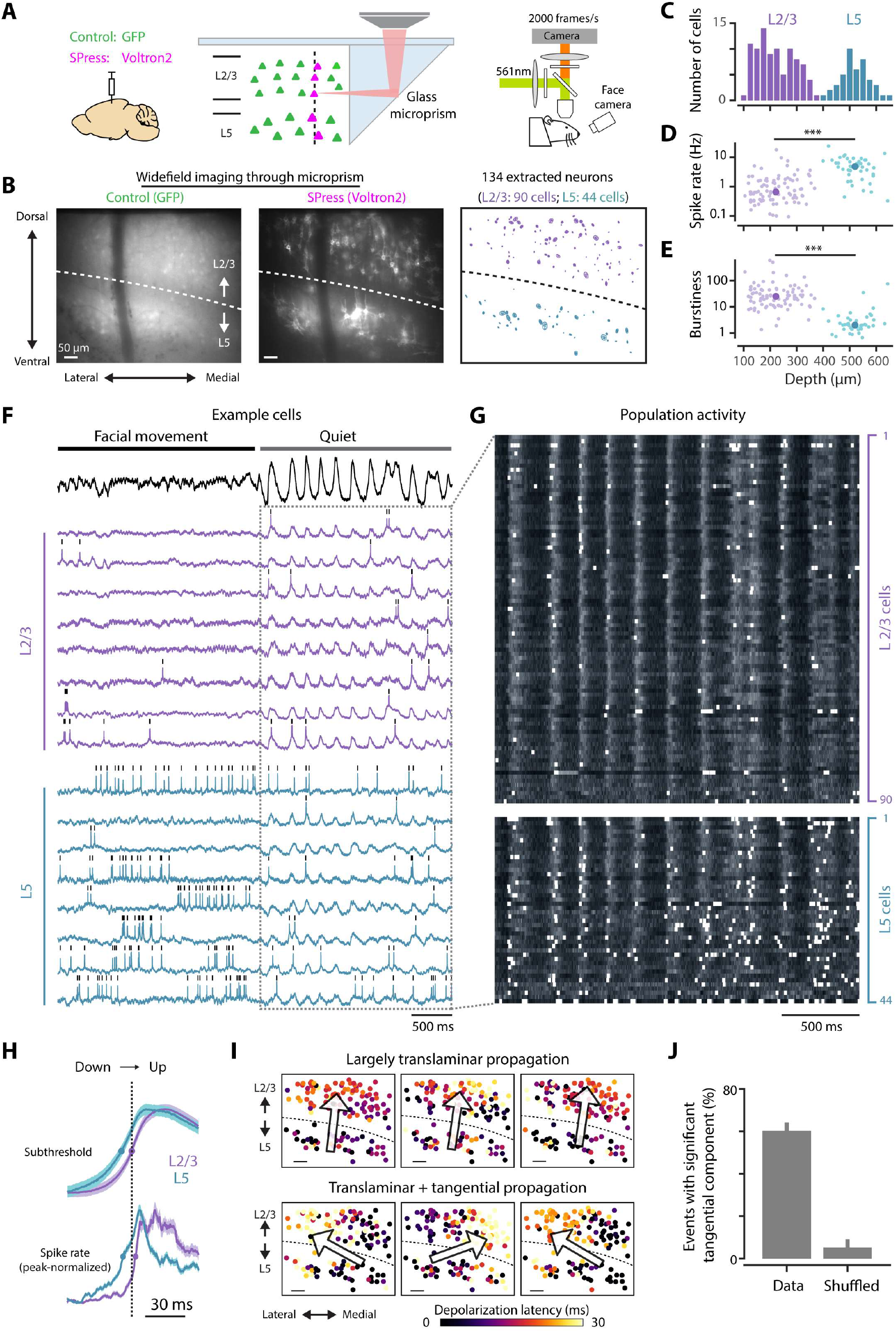
Vertical SPress reveals event-specific recruitment of cortical layers. (**A**) AAVs for SPress-mediated Voltron2 expression (magenta) and GFP control (green) were injected during prism implantation into cortex (left), followed by two-photon activation at 900 nm along a vertical plane spanning cortical layers (middle). Widefield voltage imaging was performed at 2 kHz through a glass microprism, allowing simultaneous recording from L2/3 and L5 cells. (**B**) Widefield images acquired through the implanted microprism for control GFP (left) and SPress Voltron2 (middle) in the same field of view. Right, spatial locations of extracted neurons, color-coded by layer. Dashed line indicates approximate L2/3–L5 boundary. (**C**) Depth distribution of extracted neurons, separated into L2/3 (purple) and L5 (blue). (**D and E**) Spike rate (D) and burstiness (E) of neurons as a function of depth. ***P < 0.001, two-sided t-test. (**F**) Example extracted voltage traces from L2/3 (purple) and L5 (blue) neurons during facial movement and quiet periods. Tick marks indicate detected spikes. Top, population-averaged subthreshold voltage signal. (**G**) Population activity during the quiet period in (F) across L2/3 (top) and L5 (bottom) neurons. Grayscale indicates subthreshold membrane potential; white ticks indicate action potentials. (**H**) Subthreshold voltage signal (top) and peak-normalized spike rate (bottom) of L2/3 (purple) and L5 (blue) neurons aligned to down-to-up state transitions and averaged across neurons and events. (**I**) Example events showing spatiotemporal propagation across the vertical plane. Top, events with largely translaminar propagation. Bottom, events with combined translaminar and tangential propagation. Color indicates depolarization latency for each neuron. (**J**) Fraction of events with a significant tangential propagation component in data compared with shuffled control; error bars indicate bootstrap standard errors estimated by resampling events with replacement (n = 131 events from one animal).

Neurons in the two layers exhibited distinct physiological properties (Fig. 4C-E). Spike rate (Fig. 4D) and burstiness (Fig. 4E) differed significantly between L2/3 and L5 cells, consistent with known laminar differences (*37*). At the same time, both populations exhibited pronounced behavioral-state dependence: during quiet wakefulness, neurons in both layers displayed large-amplitude, slow voltage fluctuations that were strongly reduced during facial movement (Fig. 4F and G, ‘Quiet’ vs. ‘Facial movement’). On average, L5 neurons depolarized and spiked earlier than L2/3 neurons during transitions from hyperpolarized to depolarized phases (Fig. 4H), consistent with previous reports of deep-to-superficial recruitment (*34, 38, 39*).

Population imaging across the full vertical sheet, however, revealed that this average translaminar sequence did not fully describe individual events. Latency maps of single depolarizing events showed that, while some waves propagated predominantly along the deep-to-superficial axis (Fig. 4I, top), many others combined translaminar propagation with a substantial tangential component (Fig. 4I, bottom). In this vertical-plane recording, the tangential component could favor either the medial or lateral direction and was statistically significant in ∼60% of events (Fig. 4J). Thus, cross-layer recruitment was not fully described by a fixed laminar sequence. Individual waves instead followed event-specific trajectories spanning both cortical depth and tangential position.

### Cellular-scale wavefront microstructure within propagating cortical dynamics

As cortical waves propagated, did neuronal recruitment follow a smooth wavefront, or did waves contain fine-scale organization at cellular resolution? To address this question, we fitted each event with a plane relating depolarization latency to neuronal position, defining the wavefront arrival time expected from smooth propagation (Fig. 5A). We then examined each neuron’s recruitment relative to this fitted wavefront (Fig. 5B). Small deviations indicated recruitment consistent with smooth propagation (gray cells, Fig. 5B), whereas large deviations revealed cellular-scale microstructure that a smooth wavefront could not explain (blue and red cells in Fig. 5B).

**Fig. 5.**
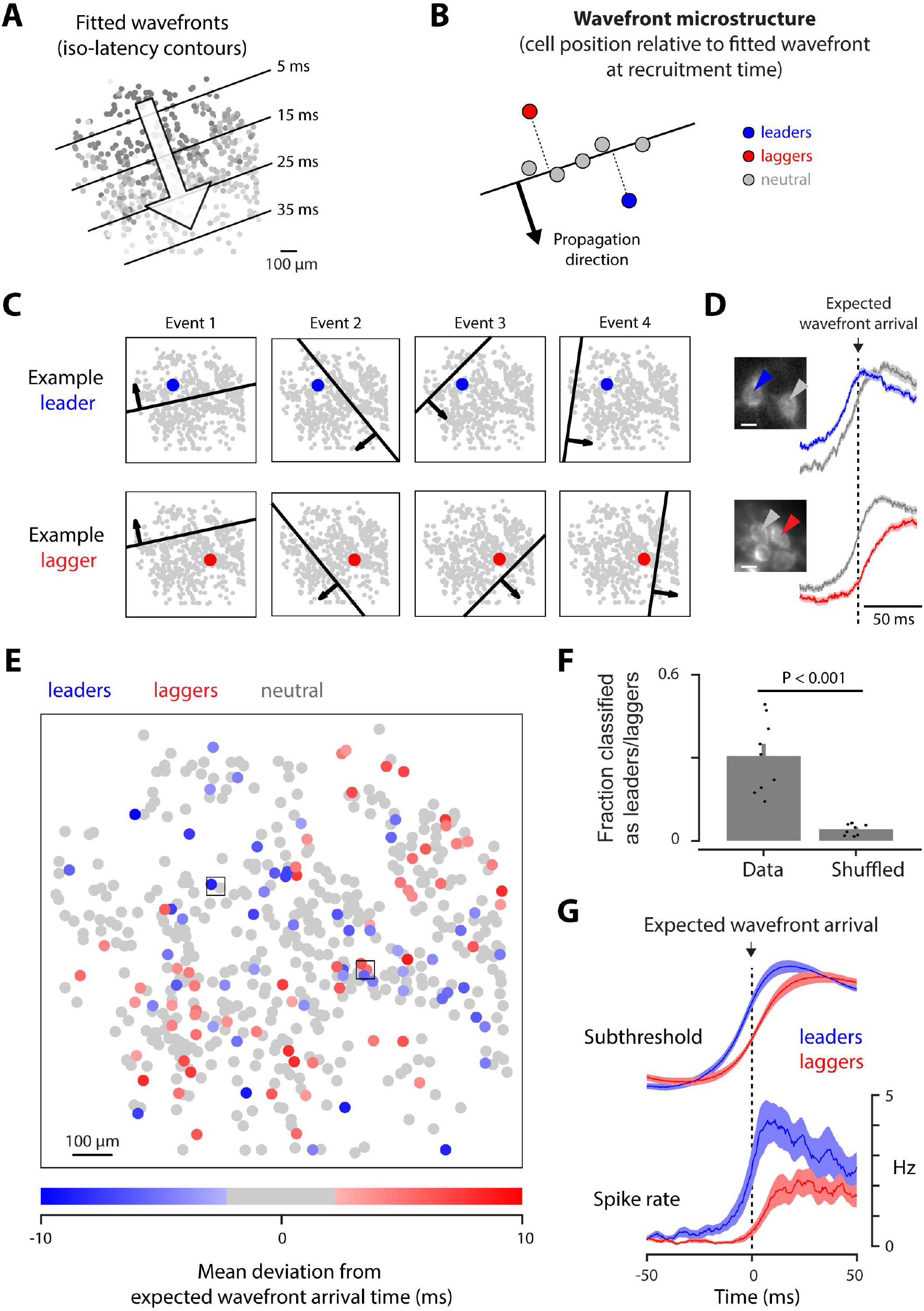
Cellular-scale wavefront microstructure within propagating cortical dynamics. (**A**) For each event, a plane was fit to neuronal depolarization latency as a function of position, defining fitted wavefronts as iso-latency contours. This fit provided the expected wavefront arrival time for each cell in each event. (**B**) Schematic illustrating wavefront microstructure. At the time of recruitment, each cell’s timing was compared with the expected arrival of the fitted wavefront. Cells recruited ahead of the fitted wavefront were defined as leaders, cells recruited behind it as laggers, and cells recruited consistently with it as wavefront-neutral. (**C**) Example leader cell and lagger cell across four events. In each event, the fitted wavefront is shown relative to the example cell at its recruitment time. (**D**) The example leader cell (blue, top) and lagger cell (red, bottom) in (C), each shown together with a nearby neuron (gray) indicated by arrowheads in the inset images. Right, average subthreshold signals across 72 events, aligned to the expected wavefront arrival time for each event. **(E**) Spatial distribution of cells colored by their mean deviation from expected wavefront arrival time across events. Cells whose mean deviation differed significantly from zero (two-sided t-test, P < 0.05) were classified as leaders or laggers; non-significant cells are shown as wavefront-neutral. Boxed cells correspond to the examples in (C) and (D). **(F)** Fraction of cells classified as leaders or laggers in the observed data and after shuffling cell identities across events. Bars show the mean, error bars indicate s.e.m., and dots represent individual animals (*n* = 9 animals). P < 0.001, two-sided paired *t*-test. **(G)** Average subthreshold signals and spike rates for leaders and laggers, aligned to the expected arrival time of the fitted wavefront at each neuron’s location. Traces were averaged across events and cells within each animal and then across animals. Shaded regions indicate s.e.m. across animals (*n* = 9 animals).

Although individual events propagated in different directions, some neurons consistently depolarized ahead of or behind the fitted wavefront (Fig. 5C). These wavefront leaders and laggers were recruited before or after the expected wavefront arrival at their locations, regardless of propagation direction (Fig. 5C and fig. S6). Within local neighborhoods, they typically depolarized earlier or later than nearby neurons (Fig. 5D). Across nine animals, 31 ± 13% of neurons (mean ± SD) were classified as leaders or laggers and were intermingled with wavefront-neutral neurons throughout the field (Fig. 5E). This fraction was substantially greater than that obtained after shuffling cell identities across events (Fig. 5F; *P* < 0.001, two-sided paired *t*-test). Leader and lagger identities were largely preserved when wavefronts were modeled using a quadratic fit to account for curvature (fig. S7), and similar local timing biases were observed in the vertical-plane microprism recording (fig. S8).

These timing biases, defined from subthreshold dynamics, also predicted neuronal spiking relative to local wavefront arrival. After aligning each neuron’s spike-rate profile to the expected wavefront arrival time at its location, leader cells spiked earlier and at higher rates than simultaneously recorded lagger cells (Fig. 5G, fig. S8). Thus, cortical voltage dynamics are not fully captured by smooth traveling waves alone. Rather, propagating waves contain reproducible cellular-scale microstructure, revealing fine-scale order within mesoscopic cortical dynamics.

## Discussion

Optical sectioning need not rely on complex optical design; it can instead be encoded genetically. By restricting fluorescence to a user-defined plane, SPress suppresses background at its source, preserving the speed and spatial coverage of widefield imaging while providing optical-sectioning-like contrast. This addresses a central barrier in voltage imaging, where kilohertz recording from densely labeled populations has required specialized instrumentation and restricted fields of view (*14*) (fig. S1). SPress enables millimeter-scale, cellular-resolution voltage imaging in dense neuronal populations with standard widefield microscopes, potentially bringing high-speed population voltage imaging within reach of many neuroscience laboratories.

The importance of this capability is not simply that more neurons can be recorded at higher speed. SPress opens an experimental regime between classical electrophysiology and mesoscale imaging: intracellular electrophysiology provides direct access to membrane voltage (*24*) but only from a small number of neurons, whereas calcium and widefield imaging (*28, 40*) capture activity across large populations but without simultaneous access to fast subthreshold voltage or spike timing from individual cells. By enabling simultaneous measurements of subthreshold voltage and spiking from hundreds of individually resolved neurons across millimeter-scale fields, SPress links cellular voltage to mesoscopic brain dynamics.

Our results suggest that large-scale voltage propagation provides a flexible, event-specific scaffold for cortical coordination, shaping when neurons depolarize, when they spike, and how layered circuits are recruited. In vertical planes, cross-layer recruitment was often accompanied by substantial lateral propagation, indicating that apparent translaminar sequences can arise as oblique intersections through broader traveling waves rather than purely vertical layer-to-layer flow. Local spike sequences and layer-to-layer timing therefore are not fixed outputs of circuit architecture alone, but depend on how each propagating event intersects the cortical sheet and columnar architecture.

Beyond these mesoscale patterns, our results reveal fine-scale cellular timing structure that smooth wavefronts alone cannot explain. As waves traveled across cortex from different directions, some neurons consistently led or lagged the local wavefront, suggesting that a cell-specific landscape shapes the timing of recruitment during propagating activity. Subthreshold waves therefore organize cortical spike timing in a manner that is simultaneously flexible and stereotyped: their trajectories reconfigure firing order at the network scale, while stable cell-specific recruitment biases shape local timing.

The same strategy could reveal additional forms of multiscale organization across brain regions and functional signals. Because SPress defines the expression plane through two-photon photoactivation, it could in principle be positioned wherever optical access can be obtained, including deep brain structures accessed through microprisms or other implantable optical interfaces. Its modular recombinase design should also be compatible with other genetically encoded functional indicators, including calcium (*41*), neurotransmitter (*42*), and neuromodulator sensors (*43, 44*), potentially allowing standard widefield microscopes to image diverse functional readouts with high speed and contrast. This capability may be particularly useful for head-mounted imaging in freely moving animals (*45*), where background fluorescence often degrades signal quality and compact hardware limits optical-sectioning options.

Future developments could further expand the capabilities of SPress. Although genetic confinement reduces out-of-focus background, tissue scattering can still blur cellular signals at greater depths. Red-shifted indicators (*46, 47*), in vivo tissue clearing (*48*), and computational demixing (*16, 49*) may mitigate this effect. A further practical consideration is the time and light exposure required for photoactivation, particularly for larger planes. Neurons within activated planes nevertheless exhibited expected physiological features, including behavior-dependent membrane-potential dynamics, Up–Down-state transitions, and layer-specific differences in firing and bursting, suggesting that gross physiological function was preserved. Ongoing efforts to improve photoactivatable recombinase efficiency (*50*) could shorten photoactivation and reduce light exposure, whereas direct plane-selective activation of fluorogenic dyes (*51*) or photoactivatable GEVIs (*52*) could eliminate the subsequent reporter-expression delay. Finally, the current implementation, like other widefield approaches, records a single optical plane. Intersectional genetics (*53*), orthogonal recombinases (*54*), and spectrally or polarity-distinct indicators (*46, 55*) could enable cell-type-specific targeting and multiple distinguishable expression planes.

Since the time of Cajal, selective labeling has been central to neuroscience, revealing neuronal structure by highlighting selected cells within complex tissue. SPress extends this principle to functional imaging, allowing activity to be isolated from a defined plane within dense tissue. More broadly, SPress reframes optical constraints as problems of biological design, opening the door to imaging strategies in which spatial selectivity is specified not only by the microscope, but also by the cells themselves.

## Acknowledgments

We thank L. Lavis for providing the JF dyes used in this study; E. Schreiter, I. Kolb and A. Abdelfattah for sharing the Voltron2 viruses; and S.-C. Lin, T. Engel, and Z. Jiang for comments on the manuscript. The authors used ChatGPT, Gemini, and Claude to assist with language editing and refinement of manuscript organization. All scientific content, citations, and final wording were reviewed and verified by the authors.

## Author contributions

Conceptualization, Methodology, Investigation, Analysis, Writing: T.-W.C. and B.-J.L.

## Data, code, and materials availability

All data needed to evaluate the conclusions in the paper are present in the paper and/or the Supplementary Materials. Source data underlying the main and supplemental figures will be deposited in a public repository before publication.

## Supplementary Materials

### Materials and Methods

#### Animals and Surgical Procedures

All procedures were performed in accordance with protocols approved by the National Yang Ming Chiao Tung University Animal Care and Use Committee. Adult C57BL/6 mice of either sex (1.5–4 months old) were used in all experiments. Mice were anesthetized with 0.8–1.5% isoflurane and maintained on a heating pad throughout surgery. A circular craniotomy (3 mm in diameter) was made over the left cerebral cortex, targeting regions with minimal surface vasculature. Across animals, the center of the craniotomy ranged from anteroposterior +1 to −4.0 mm and mediolateral 2.0 to 2.7 mm relative to bregma.

To enable Selective Plane Expression, we injected an AAV mixture comprising three components: (1) a photoactivatable recombinase, (2) a Cre-dependent fluorescent reporter (tdTomato) or voltage indicator (Voltron2-ST), and (3) a constitutive fluorescent control (GFP). Detailed descriptions of viral constructs, sources, and final concentrations are provided in Table S1. Viruses were mixed and diluted to final concentration in PBS before injection, and delivered at 5–10 sites per animal (60–80 nl per site), with injection sites spaced by ∼500 μm and targeted to a depth of ∼350 μm below the cortical surface. Injections were performed at a rate of 1 nl s⁻¹. After injection, an imaging window consisting of a 3-mm-diameter glass coverslip attached to a stainless steel cannula (3 mm diameter, 0.7 mm height) was placed on the cortex and secured to the skull using Super-Bond C&B (Sun Medical). A titanium head bar was glued to the skull for head fixation during imaging.

Microprism implantation was performed using a procedure similar to that described by Andermann et al. (2013). Briefly, a prism assembly was prepared by gluing a 1.0-mm microprism with an enhanced aluminum-coated reflective surface (MPCH-1.0, Tower Optical) to a 3.0-mm-diameter glass coverslip using optical adhesive (NOA-81, Norland). A circular craniotomy (3 mm diameter) was made over the left cerebral cortex, followed by viral injection at three sites along the mediolateral axis, spaced approximately 400 μm apart. At each site, injections were performed at two depths (350 and 700 μm below the cortical surface), with 60 nl delivered at each depth. A 1.2-mm-long incision, parallel to the injection plane, was then made into the cortex to a depth of approximately 1 mm using a dissecting knife (FST, 10055-12). The prism assembly was held with a vacuum suction needle mounted on a three-axis micromanipulator, slowly inserted into the precut slit, and secured to the skull using Super-Bond C&B.

After surgery, the imaging window was covered with a custom 3D-printed plug and sealed with Kwik-Cast (World Precision Instruments) to protect it from light exposure.

#### Two-Photon Activation of Selective Plane Expression

Selective Plane Expression was induced 2–3 weeks after surgery by two-photon activation of the photoactivatable recombinase at 900 nm. Mice were placed in a tube and head-fixed under the microscope, and the light-protection plug covering the imaging window was carefully removed with forceps. All procedures were performed under minimal ambient light using only red-light illumination to minimize unintended background recombination.

Imaging and activation were performed using a two-photon microscope equipped with a Ti:sapphire laser (Chameleon Ultra II, Coherent) and a Nikon 16× 0.8 NA objective. A suitable field of view located 100–200 μm below the cortical surface was first identified by imaging GFP fluorescence with 1000-nm excitation, which did not induce recombination (20). The excitation wavelength was then switched to 900 nm for activation. Activation was performed by raster scanning the laser focus with a resonant galvo system (RESSCAN, Sutter Instrument) at 100–150 mW, depending on the depth of the target plane. Scanning was performed at 512 × 512 pixels and 30 Hz for 30 min for a 300 × 300 μm field or 210 min for a 1000 × 1000 μm field. For microprism experiments, a 750 × 750 μm field was scanned for 270 min at 100 mW. After induction, the imaging window was re-covered with the 3D-printed plug, sealed with Kwik-Cast, and the animal was returned to its home cage.

#### Voltage Imaging

Voltage imaging was performed 2–3 weeks after SPress induction. For fluorescent labeling of Voltron2-ST, 100 nmol JF_552_-HaloTag ligand was dissolved in 20 μl DMSO (Sigma), diluted first in 20 μl Pluronic F-127 [20% (wt/vol) in DMSO; P3000MP, Invitrogen], and then in 80 μl PBS. The dye solution was administered by retro-orbital injection using a 30-gauge needle.

Imaging sessions were performed 12–24 h after dye injection on a widefield microscope built on a Thorlabs Cerna platform. A 561-nm laser (Jive 561, Cobolt) was expanded and projected onto the sample at an intensity of 25–50 mW mm⁻² to excite Voltron2-labeled neurons. Fluorescence was collected through a 16× 0.8-NA objective (Nikon), separated from excitation light by a dichroic mirror (570lpxr, Chroma) and an emission filter (ET605/70, Chroma), and directed through a dual camera port (Y-IDP, Nikon). For GFP imaging, samples were illuminated with a blue LED (M470L3, Thorlabs) and imaged using a standard GFP filter set (49002 ET, Chroma).

High-resolution widefield images of the SPress-labeled region were first acquired for both GFP and Voltron2 channels using an sCMOS camera (Zyla 4.2, Andor) at 2048 × 2048 pixels (0.58 μm per pixel). For voltage imaging, Voltron2 fluorescence was directed to the front port of the Y-IDP and imaged with a high-speed CMOS camera (DaVinci-1K, RedShirt) at 2000 frames per second. A 50-mm camera lens (NIKKOR 50 mm f/1.2, Nikon) was used as the tube lens in this optical path, producing an effective magnification of ∼4×. Images were acquired using Turbo-SM64 software (SciMeasure) for 3 min (360,000 frames) at 190 × 160 pixels (7.5 μm per pixel), covering an area of approximately 1.4 × 1.2 mm. During imaging, animals were fully awake and head-fixed, with free movement of whiskers and limbs.

#### Behavioral Imaging

Animal behavior was monitored during voltage imaging using a video camera. The animal’s face was illuminated with a 685-nm LED (JET-685-05, Roithner) through an excitation filter (AT690/50, Chroma) and imaged through a 50-mm lens (TC5028-3MP, Tokina) onto a CCD camera (BFLY-U3-05S2M, FLIR) positioned ∼20 cm from the face. Images were acquired at 808 × 608 pixels and 25 Hz and synchronized to voltage imaging via TTL pulses from Turbo-SM64.

#### Two-Photon z-Stack Imaging

After widefield voltage imaging, animals were transferred to a two-photon microscope for structural imaging. The SPress-labeled region was identified, and a two-photon z-stack was acquired at 2-μm axial spacing and 512 × 512 pixel resolution. Voltron2-JF_552_ and tdTomato fluorescence were imaged using 850 nm and 1000 nm excitation, respectively, and collected through an emission filter (FF01-609/57-25, Semrock). GFP fluorescence was also excited by these wavelengths and acquired simultaneously with the red channel.

#### Motion estimation and correction

Lateral brain motion was estimated frame-by-frame using a gradient-based least-squares method. A reference image *f*_0_(x, y) was defined as the mean of the first 2000 images of each recording. Assuming small translational displacements (Δ*x*, Δ*y*), each frame was approximated as 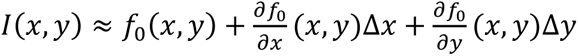. The spatial gradient images, 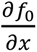 *and* 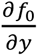, were computed from the reference image using centered finite differences, and (Δ*x*, Δ*y*) was estimated for each frame by least-squares fitting to this linearized model. Each frame was then translated by the negative of its estimated displacement to align the movie to the reference image.

#### Neuron identification and signal extraction

Neurons and their activity traces were identified using activity localization imaging (ALI) (16), which localizes fluorescence transients associated with individual action potentials (APs) to visualize and separate nearby neurons. Briefly, motion-corrected movies were first high-pass filtered pixelwise by subtracting a slow baseline estimated with a 10-ms median filter to isolate rapid spiking signals. Because Voltron fluorescence decreases with depolarization, the filtered movie was inverted so that APs appeared as positive events. Candidate APs were detected by spatial smoothing of the high-pass-filtered movie with a Gaussian kernel approximately matched to neuronal size, pixelwise noise normalization, thresholding, and grouping suprathreshold voxels into spatiotemporally connected events. For each event, spike position was refined to subpixel precision by centroid-based localization. The refined locations of all APs were then accumulated into an AP-density map, from which cluster centers corresponding to individual neurons were identified as local maxima. Low-firing-rate clusters missed by the automated procedure were added manually when necessary. Each AP was assigned to the nearest cluster, and neuronal footprints were estimated by averaging spike images within each cluster after excluding synchronous spikes from other clusters that could contaminate the average. Voltage traces were then extracted by modeling the movie as a linear combination of these footprints and solving for the time-varying coefficients using non-negative least-squares regression. Spike times, footprints, and traces were further refined by repeating spike detection, footprint estimation, and trace extraction twice. To recover full voltage traces containing both spiking and subthreshold dynamics, the same footprints identified from the high-pass-filtered movie were used for regression on the original movie. A slow baseline, estimated for each trace using a 1-s moving mean, was then subtracted to remove photobleaching and other slow fluctuations. Extracted voltage traces were normalized to the spike height of the same cell. For subthreshold analysis, spike-related signals were removed from each trace by excluding samples within a −4 to +7 frame window around each detected spike, followed by linear interpolation across the excluded interval using the nearest samples outside the window.

#### Spike-rate and burstiness analysis

Spike rate was calculated for each neuron as the number of detected spikes divided by the duration of the recording period. Burstiness was quantified using an autocorrelogram-based index (37). For each neuron, all pairwise spike-time differences were computed, histogrammed at the imaging frame interval, and normalized by the total number of spikes, yielding the probability of observing another spike at each time lag relative to a reference spike. Burstiness was defined as the mean spike probability at positive lags of 7–28 frames after each spike, corresponding to 3.5–14 ms at 2000 frames s⁻¹, divided by the neuron’s overall mean spike probability per frame. This dimensionless index quantifies the excess probability of short-latency spike recurrence relative to the neuron’s baseline firing probability.

#### Behavioral State Classification

To detect periods of facial movement and quiet immobility, movement strength was estimated for each frame of the behavioral movie as the sum of squared pixel-value differences between consecutive frames. Low and high thresholds were defined as the 50th and 80th percentiles of movement strength over the recording period. Episodes of facial movement were identified as continuous periods lasting at least 500 ms during which all samples exceeded the low threshold and at least one sample exceeded the high threshold. Episodes of quiet immobility were identified using the same procedure applied to the inverted movement-strength trace.

#### Identification of large depolarization events

Large, synchronized depolarizing events were identified from the population-averaged subthreshold signal, computed by averaging the spike-removed subthreshold traces of all neurons. Prominent peaks were detected using findpeaks in MATLAB with the minimum peak prominence set to 1 standard deviation of the population-averaged subthreshold trace; valleys were identified by applying the same procedure to the inverted trace. Large depolarizing events were then defined as peaks immediately following a valley, with a valley-to-peak amplitude greater than 4 standard deviations of the population-averaged subthreshold trace. For each such event, the Down-to-Up transition time was defined as the first sample at which the signal crossed 50% of the valley-to-peak amplitude. Subthreshold traces and spike counts of individual neurons were aligned to this transition time and averaged across cells and events.

#### Depolarization latency and propagation

For each large depolarizing event, depolarization latency of individual neurons was defined as the time point of steepest rise in the spike-removed subthreshold trace. For each neuron, a ±100-frame window centered on the population-defined event time was extracted, and the temporal derivative was computed using a symmetric difference operator with a ±10-frame offset. The derivative trace was then smoothed with a Gaussian filter (window, 40 frames), and latency was taken as the time point of its maximum within the analysis window.

For each depolarizing event, propagation was quantified by fitting a plane to the depolarization latencies of individual neurons as a function of their spatial positions (fitlm, MATLAB). The fitted coefficients, which define the spatial gradient of latency, were used to determine propagation direction. Statistical significance was assessed from the p-values of these coefficients. Events with significant lateral propagation were defined as those with a significant regression coefficient in the x direction (P < 0.05).

To estimate the chance level for detecting significant lateral propagation, depolarization latencies were randomly permuted across neurons independently for each event, while neuronal positions were kept fixed. This shuffle preserved the latency distribution within each event but disrupted the spatial relationship between latency and cell position. Planar regression was then repeated on the shuffled data, and the fraction of shuffled events with a significant x-gradient was used as the chance level. Error bars were estimated by bootstrap resampling events with replacement.

To compare neuronal activity on the upstream and downstream sides of each propagating event, neurons were grouped by their spatial position along the event-specific propagation axis. For each event, the fitted spatial latency gradient defined the propagation axis, with the gradient pointing toward increasing depolarization latency. The position of each neuron was projected onto this axis. Neurons with projected positions below the median were classified as upstream, corresponding to the earlier-arriving side of the fitted wavefront, whereas neurons with projected positions above the median were classified as downstream. Upstream and downstream groups were defined independently for each event. Spike-removed subthreshold traces and spike-rate traces were aligned to the population-defined Down-to-Up transition time and averaged separately for upstream and downstream neurons across cells and events.

#### Wavefront microstructure analysis

To quantify how well the event-specific propagation model explained the spatial pattern of depolarization latency, we compared measured latencies with latencies predicted from planar wavefront fits. For each large depolarizing event, a plane was fitted to the depolarization latencies of all neurons as a function of their spatial positions, as described above. The fitted plane provided the expected wavefront arrival time of each neuron based on the large-scale propagation geometry of that event. For each neuron and event, residual latency was calculated as the measured depolarization latency minus the latency predicted by the fitted plane. Negative residuals indicated that a neuron depolarized earlier than expected from the fitted wavefront, whereas positive residuals indicated later-than-expected depolarization.

To identify neurons with consistent timing biases relative to the event-specific wavefronts, residual latencies were collected across events for each neuron. Neurons with a mean residual significantly different from zero were identified using a two-sided one-sample t-test across events. Neurons with significantly negative mean residuals were classified as leader cells, whereas neurons with significantly positive mean residuals were classified as lagger cells.

To relate residual subthreshold timing bias to voltage dynamics and spike output, voltage traces and spike trains were aligned to the expected local timing of each event. For each neuron and event, the expected local timing was defined as the depolarization latency predicted by the event-specific planar wavefront fit. Spike-removed subthreshold traces and detected spike trains were then aligned to this predicted local timing and averaged across events for leader and lagger cells. This alignment removed timing differences expected from large-scale propagation, allowing cellular voltage dynamics and spiking to be compared relative to the local wavefront. Thus, time zero in this analysis represents the time at which the fitted wavefront was expected to reach each neuron, rather than the global population event time.

**Fig. S1.**
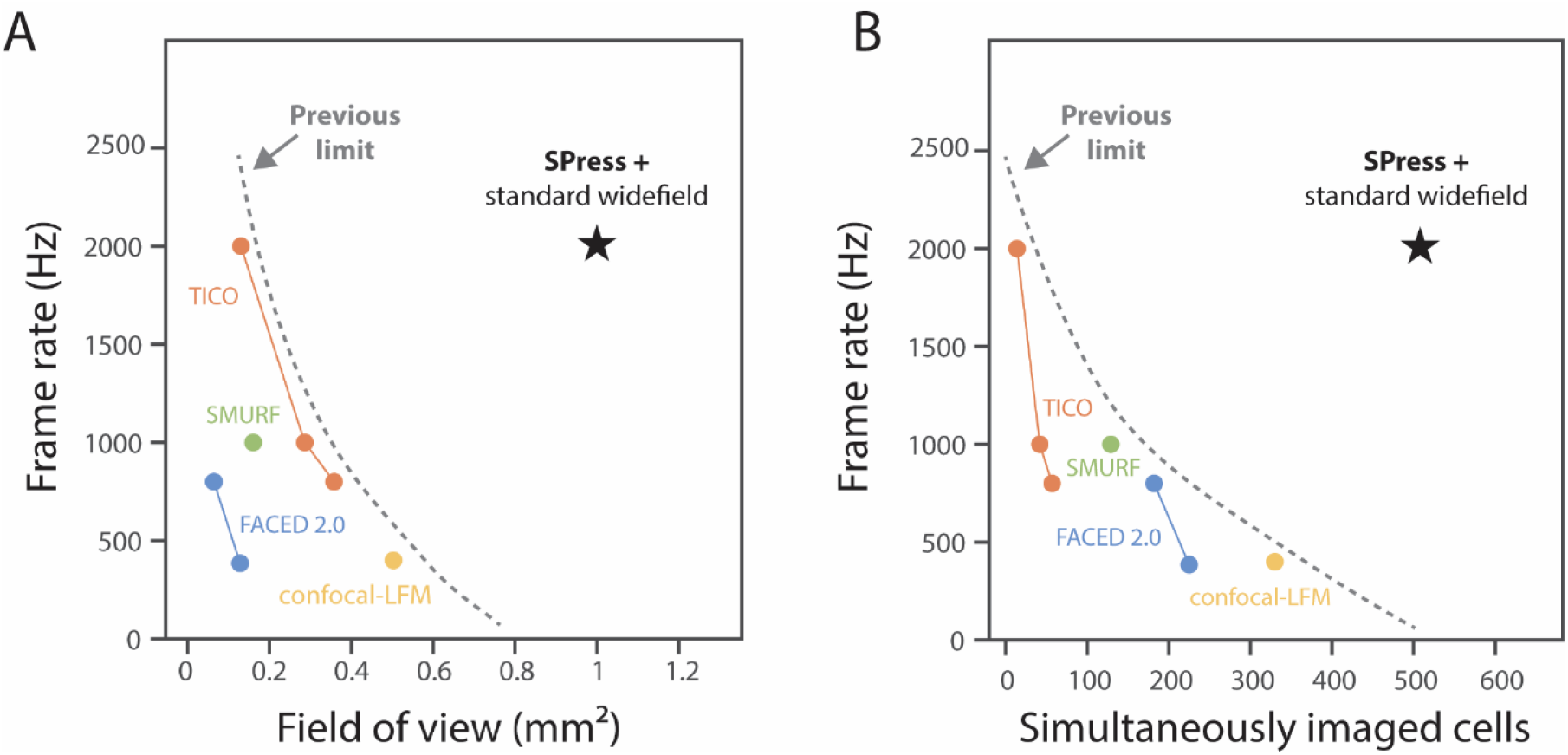
SPress extends the speed–scale regime of voltage imaging without specialized optical-sectioning hardware. (**A**) Frame rate versus field of view for representative optical-sectioning approaches used for voltage imaging. The dashed curve indicates the approximate performance frontier of previous high-speed optical-sectioning approaches, which require specialized optical-sectioning hardware during recording. The star marks SPress, in which optical-sectioning-like contrast is generated by a one-time expression-patterning step, allowing multi-kilohertz voltage imaging over a ∼1-mm^2^ field of view using a standard widefield microscope. (**B**) Frame rate versus the number of simultaneously imaged neurons for the same methods. SPress combines kilohertz imaging with simultaneous recording from hundreds of neurons.

**Fig. S2.**
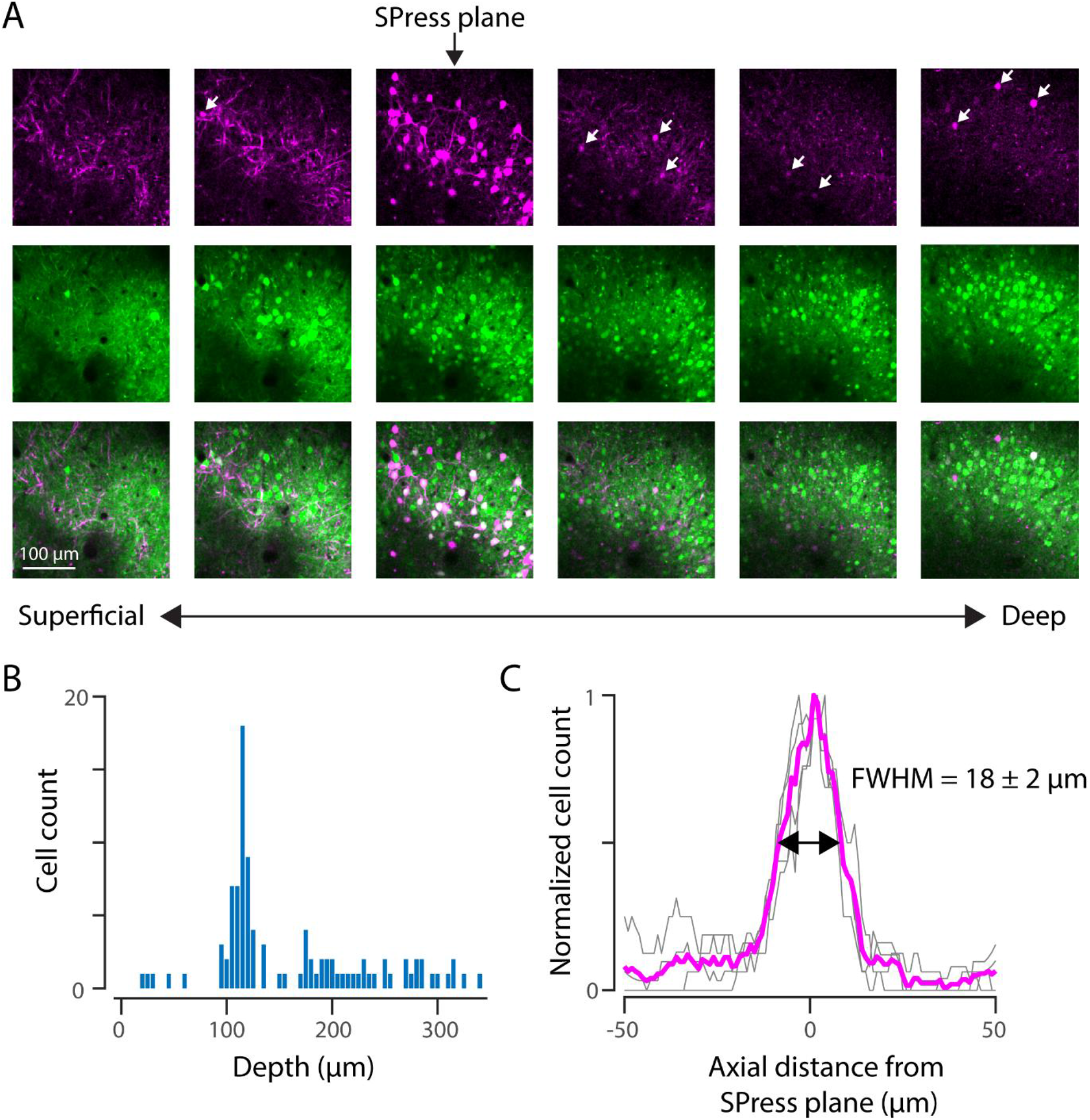
SPress labeled cell number as a function of depth. (**A**) Two-photon images of SPress-labeled neurons (magenta) and control GFP fluorescence (green) at increasing depths from superficial to deep cortex. Arrows indicate labeled neurons outside the SPress plane. Bottom row shows merged images. (**B**) Distribution of labeled neurons as a function of depth. Each bar represents the number of labeled cells within a 5-μm depth bin. (**C**) Axial profile of labeled cell density, aligned to the SPress activation plane (0 μm). Magenta line shows the mean normalized cell count across experiments; gray lines indicate individual datasets.

**Fig. S3.**
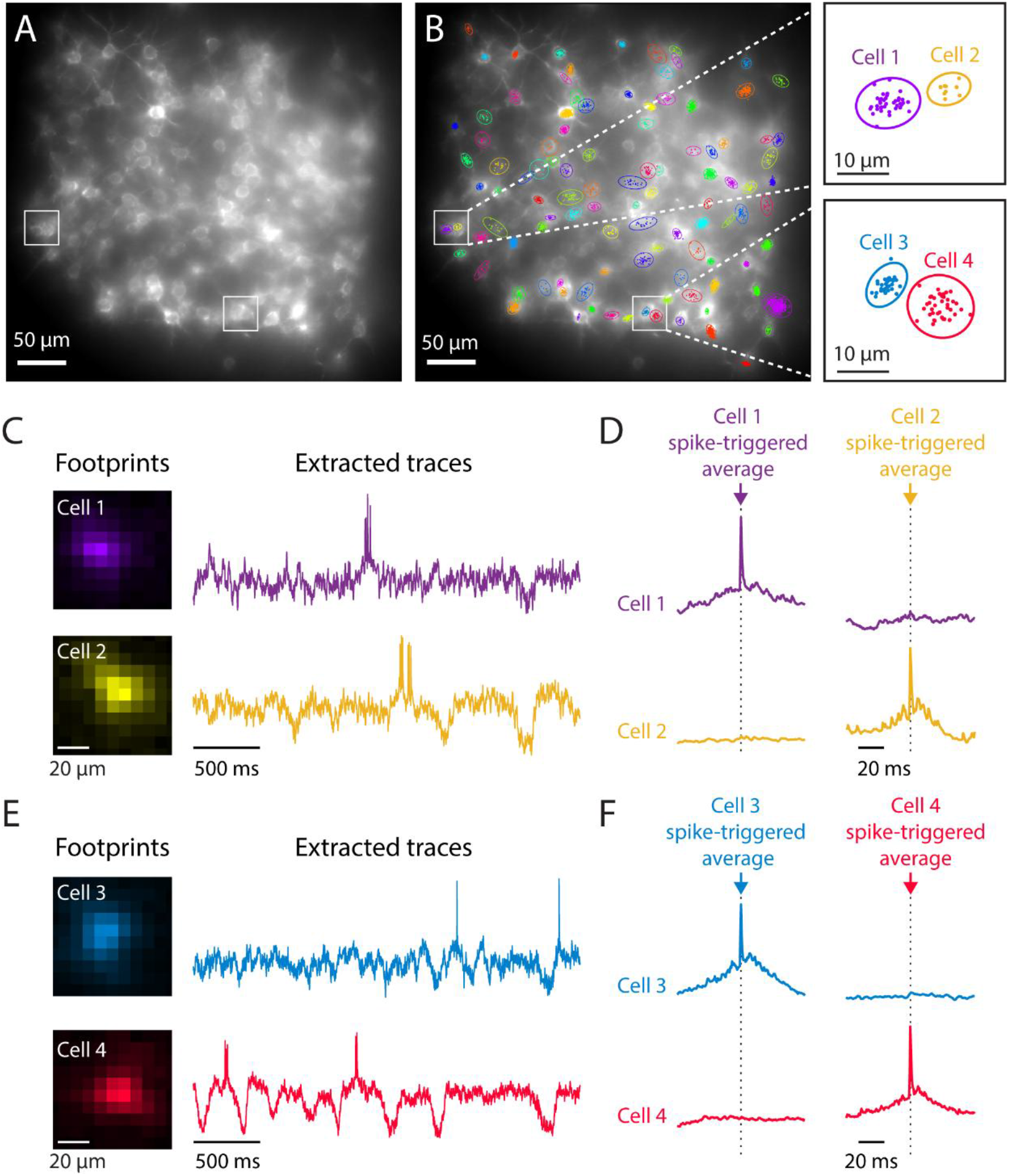
Separation of voltage signals from closely spaced neurons. (**A**) Widefield fluorescence image of an SPress-labeled neuronal plane. White boxes indicate pairs of closely spaced neurons shown in subsequent panels. (**B**) Same field of view with localized spike events overlaid. Colored outlines indicate distinct spike clusters. Dashed lines indicate zoomed regions shown at right. Right, magnified views of localized spike events and corresponding clusters. (**C**) Spatial footprints and extracted voltage traces for Cell 1 and Cell 2. (**D**) Spike-triggered averages aligned to spikes from Cell 1 or Cell 2. (**E**) Spatial footprints and extracted voltage traces for Cell 3 and Cell 4. (**F**) Spike-triggered averages aligned to spikes from Cell 3 or Cell 4.

**Fig. S4.**
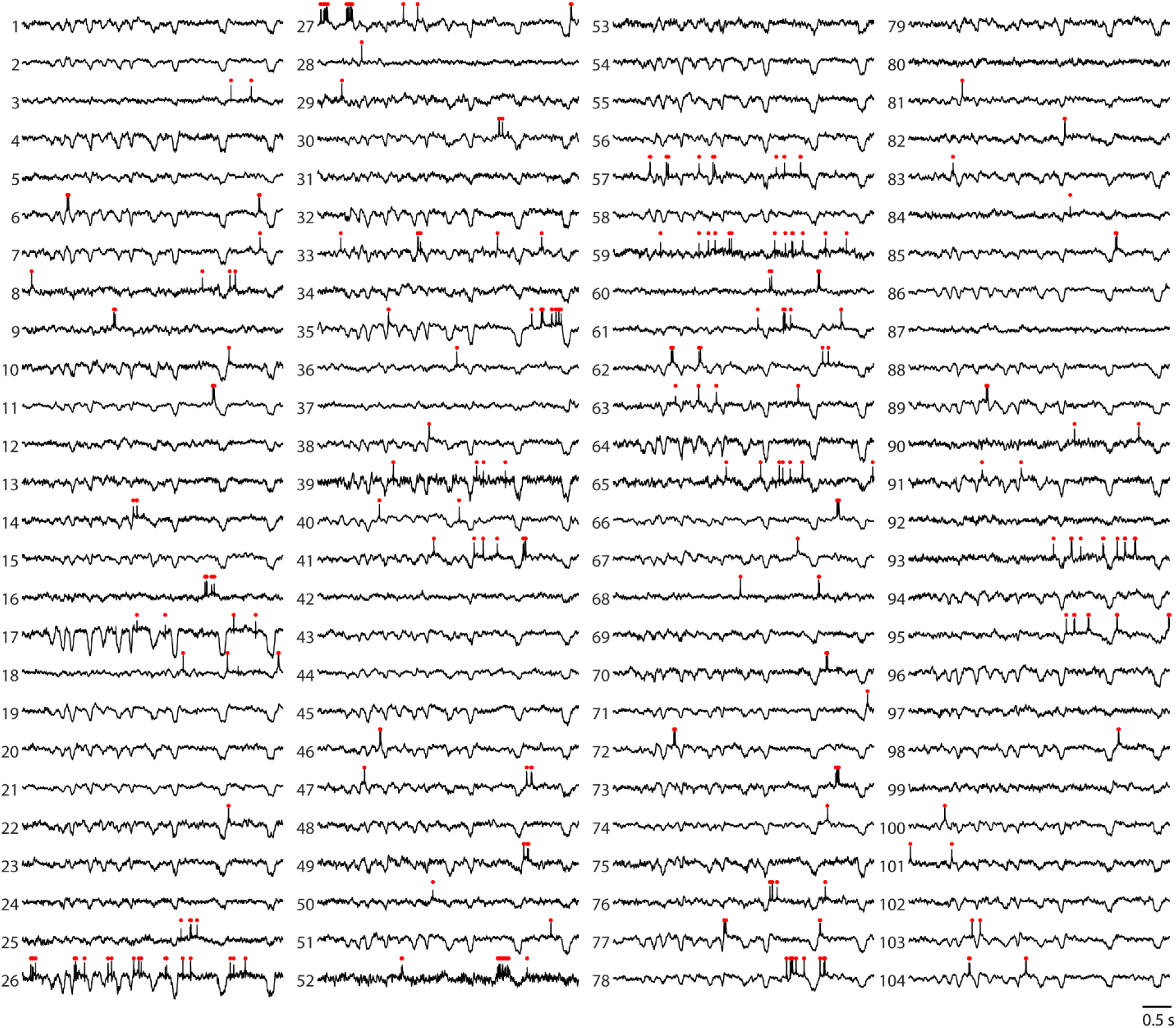
Extracted voltage traces from the example field of view. Voltage traces extracted from the 104 neurons shown in Fig. 2, displayed over a representative time window. These traces include both subthreshold voltage fluctuations and spiking activity. Red ticks indicate detected action potentials.

**Fig. S5.**
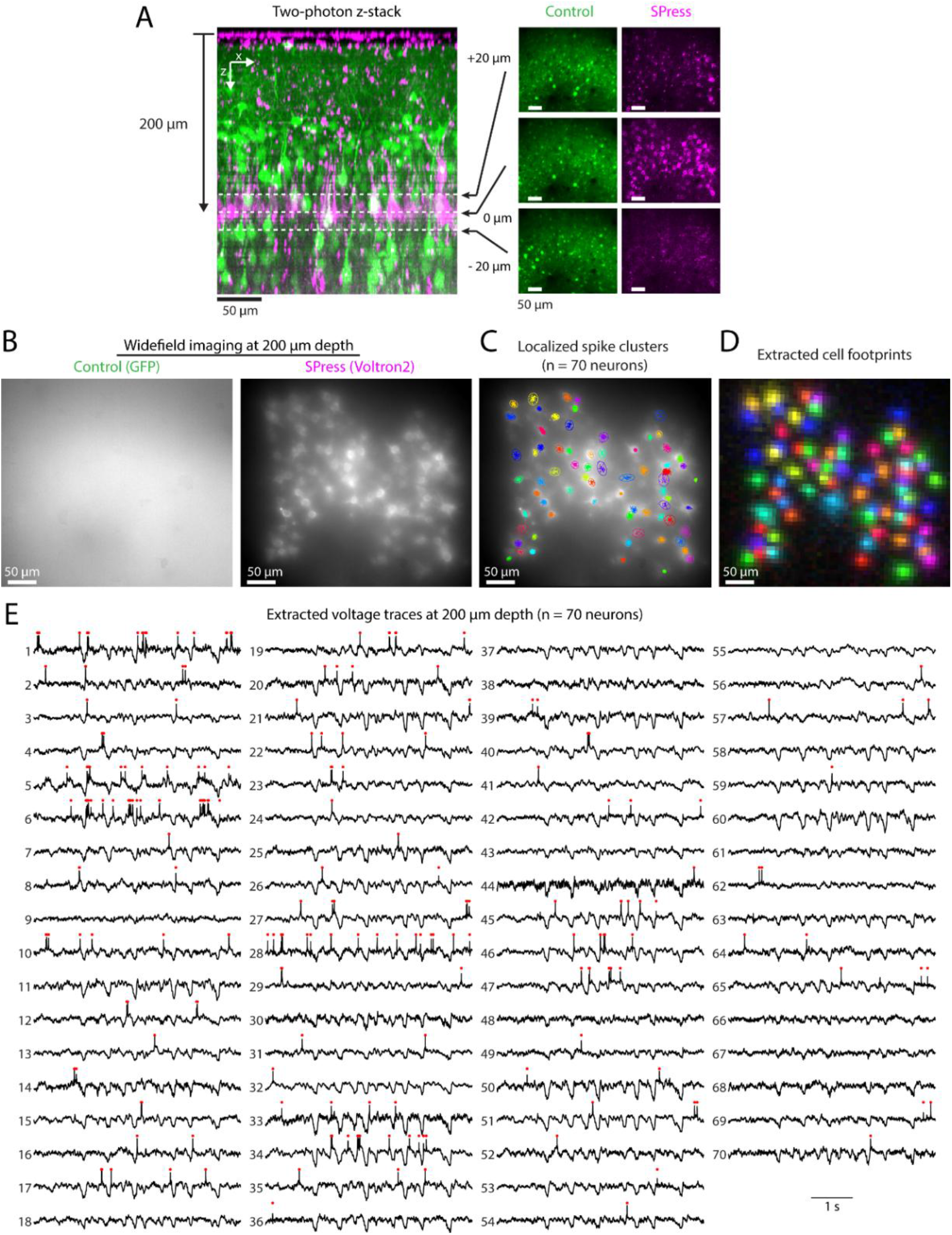
Example of SPress voltage imaging at 200 μm depth. (**A**) Two-photon z-stack showing the SPress-labeled plane located 200 μm below the cortical surface. Right, two-photon images acquired at the indicated depths relative to the labeled plane for control GFP and SPress. (**B**) Widefield fluorescence images acquired at 200 μm depth for control GFP and SPress (Voltron2). (**C**) Localized spike clusters identified from the SPress field of view, corresponding to 70 extracted neurons. (**D**) Extracted cell footprints for the same neurons. (**E**) Extracted voltage traces from the 70 neurons shown in (C, D), displayed over a representative time window. Red ticks indicate detected action potentials.

**Fig. S6.**
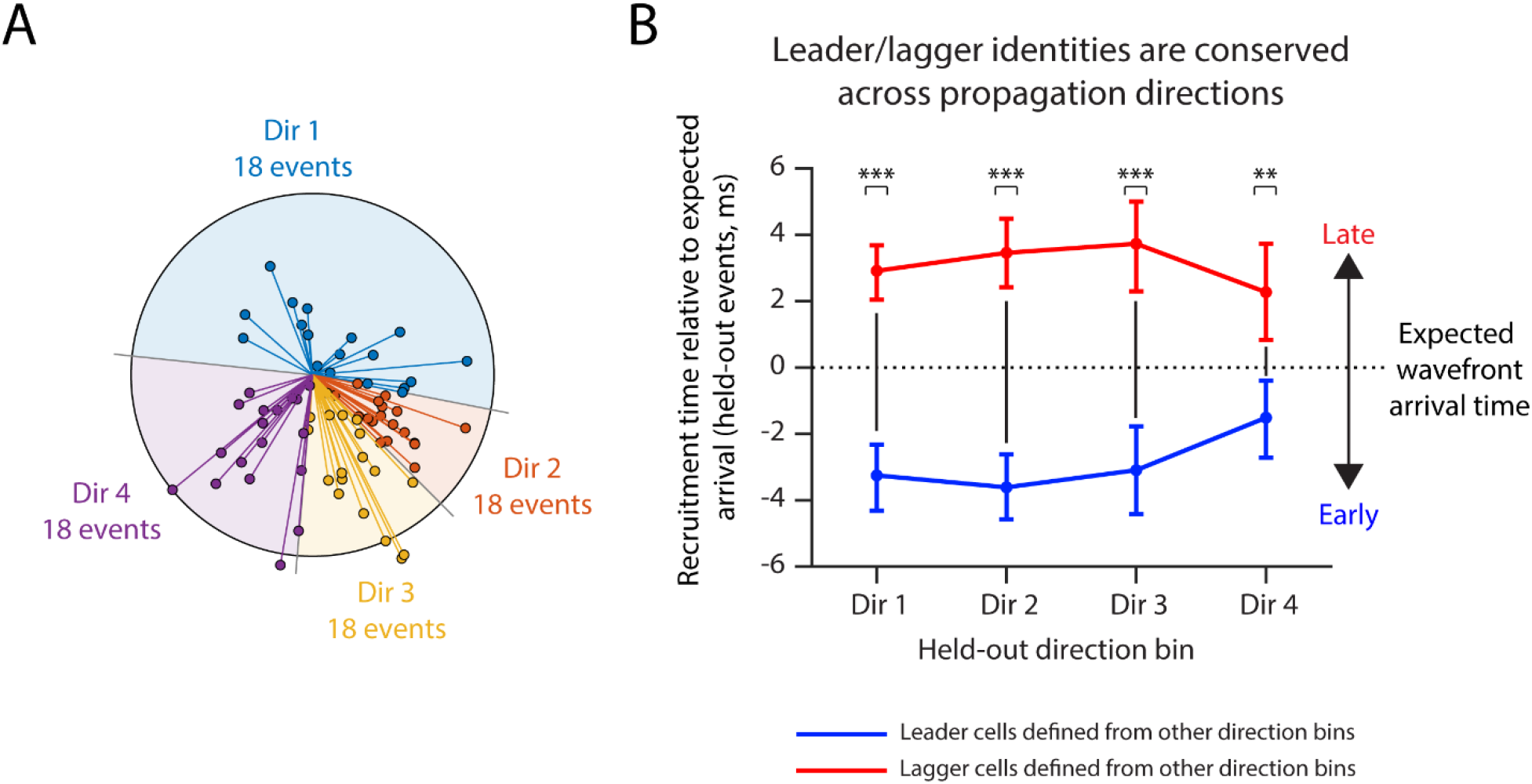
Leader and lagger identities are conserved across propagation directions. (**A**) Propagation directions of individual depolarizing events, estimated from the spatial latency gradient of the planar wavefront fit. Events were divided into four direction bins. (**B**) Leave-one-direction-bin-out analysis. For each held-out direction bin, leader and lagger cells were defined using events from the other three bins and evaluated in the held-out events. Recruitment timing is shown relative to the expected wavefront arrival time, calculated as observed recruitment time minus the arrival time predicted by the fitted wavefront. Leader cells retained earlier-than-expected recruitment, whereas lagger cells retained later-than-expected recruitment across all held-out direction bins. Error bars indicate 95% bootstrap confidence intervals. Stars indicate paired comparisons between leader and lagger recruitment timing within each held-out direction bin (***, P < 0.001; **, P < 0.01; two-sided paired t-test).

**Fig. S7.**
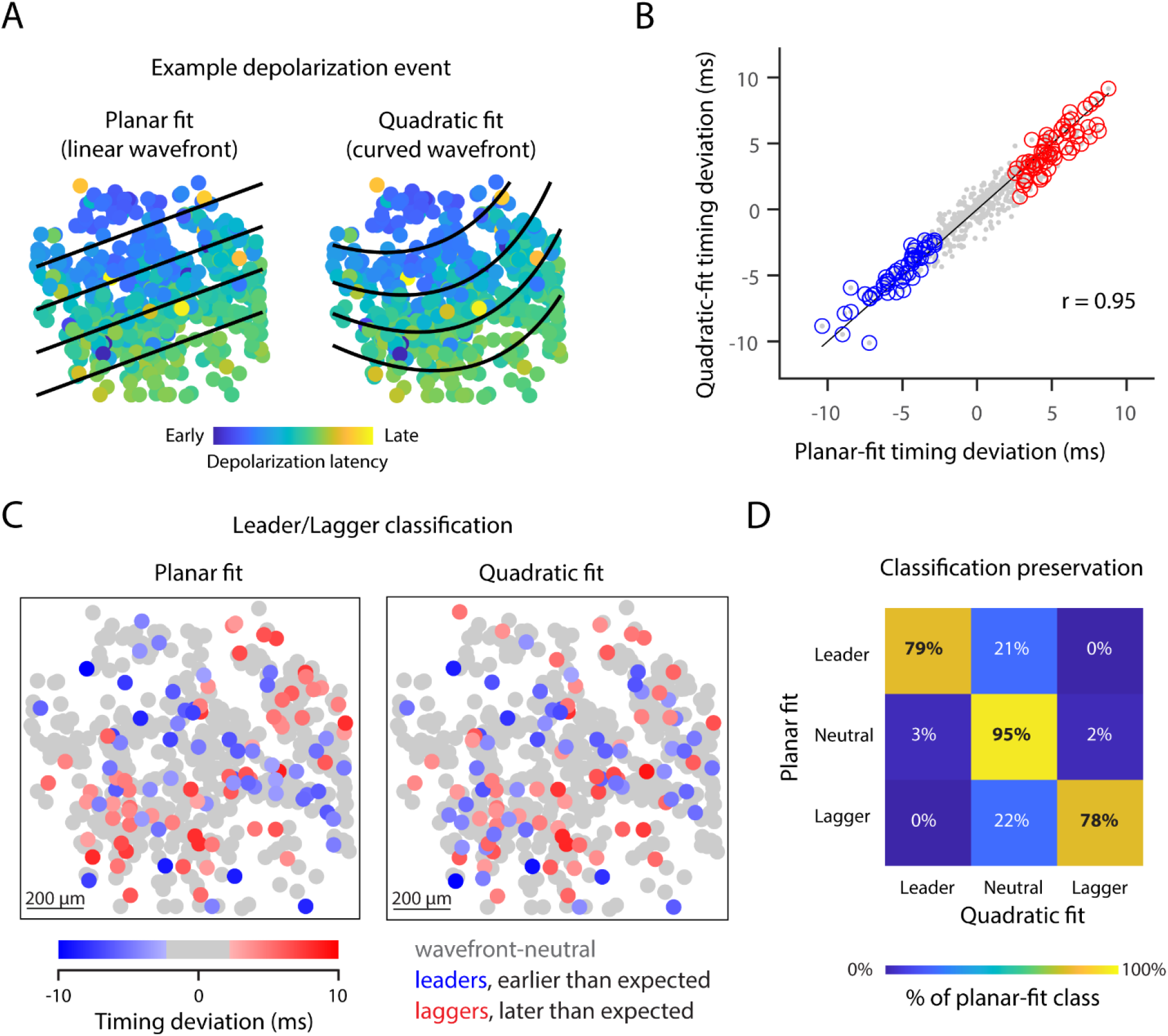
Leader and lagger identities are preserved under a curved wavefront model. (**A**) Example depolarization event fitted with planar and quadratic wavefront models. Black lines indicate predicted wavefronts at successive times. (**B**) Cell-by-cell comparison of mean timing deviation computed using planar and quadratic fits. Timing deviation was defined as the difference between observed depolarization latency and expected wavefront arrival time predicted by the fitted model, averaged across events. Each point represents one neuron and is colored by planar-fit classification. Black line, unity. (**C**) Spatial maps of leader, neutral, and lagger classifications obtained using planar and quadratic fits. (**D**) Row-normalized confusion matrix comparing classifications assigned by planar and quadratic fits. Percentages indicate the fraction of cells in each planar-fit class assigned to each quadratic-fit class.

**Fig. S8.**
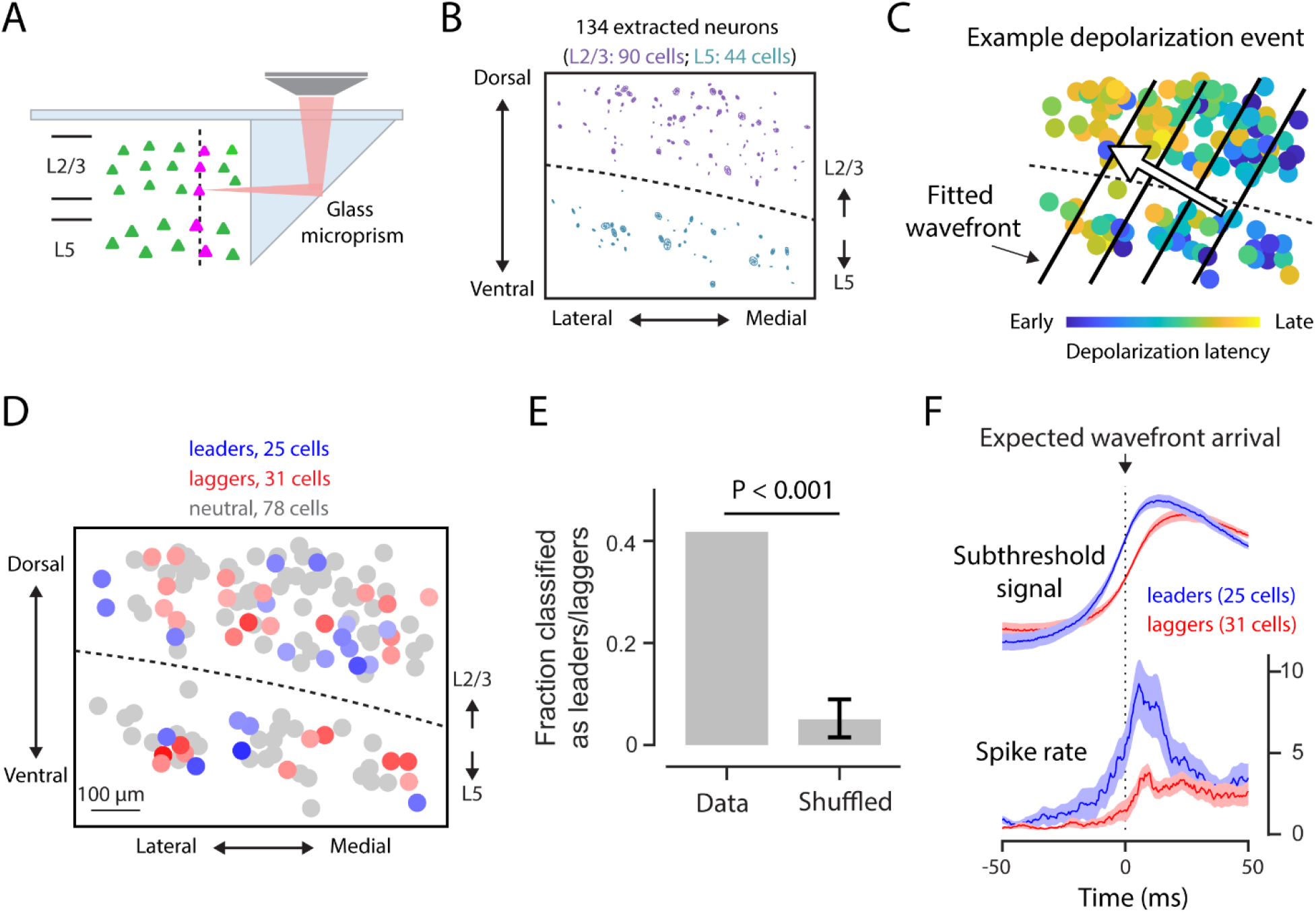
Wavefront leaders and laggers in vertical-plane microprism recordings. (**A**) Schematic of vertical-plane SPress imaging through a glass microprism. A thin expression plane was patterned across cortical depth, spanning L2/3 and L5. (**B**) Spatial distribution of 134 extracted neurons in the vertical imaging plane. Dashed line indicates the approximate boundary between L2/3 and L5. (**C**) Example depolarization event in the vertical plane. Neurons are colored by depolarization latency. Black lines indicate fitted wavefronts expected from smooth propagation. (**D**) Spatial distribution of wavefront leaders, laggers, and neutral neurons. (**E**) Fraction of neurons classified as leaders or laggers in the data compared with cell-identity shuffles. Error bar indicates the 2.5–97.5 percentile range across shuffled datasets. (**F**) Average subthreshold signals and spike rates for leaders and laggers aligned to the expected wavefront arrival time at each neuron’s location.

**Table S1.** Viral constructs used in this study.

| **Category** | **Construct** | **Source** | **Final concentration (GC/mL)** |
|---|---|---|---|
| Photoactivatable recombinases | AAV1-EF1a-iCreV | Addgene (#140135) | 2.0 × 10<sup>12</sup> |
|  | AAV1-CaMKII-iCreV | VecGene Biotechnology, Taiwan | 2.4 × 10<sup>12</sup> |
| Cre-dependent reporters | AAV1-CAG-FLEX-tdTomato | UPenn Vector Core | 6.0 × 10<sup>12</sup> |
|  | AAV1-CAG-FLEX-Voltron2-ST | Gift from Dr. Eric Schreiter (Janelia) | 2.0 × 10<sup>12</sup> |
| Control | AAV1-CB7-EGFP | UPenn Vector Core | 4.0 × 10<sup>11</sup> |

**Movie S1. Two-photon z-stack through an SPress-labeled cortical plane.**

Sequential two-photon images acquired from superficial to deeper cortical depths show SPress-mediated Voltron2 labeling (magenta) together with constitutively expressed GFP control labeling (green). Voltron2-labeled neuronal somata are concentrated within a narrow axial plane, whereas GFP fluorescence extends broadly across depth.

**Movie S2. Two-photon z-stack through a deeper SPress-labeled cortical plane.** Sequential two-photon images show an SPress-mediated Voltron2 expression plane located approximately 200 µm below the cortical surface. Voltron2-labeled neurons (magenta) are confined to a narrow plane at depth, whereas constitutively expressed GFP control fluorescence (green) remains broadly distributed across the z-stack.

**Movie S3. Z-stack comparison of SPress Voltron2 labeling and GFP control.**

Z-stack through the same cortical region showing cellular Voltron2 labeling confined to the SPress plane, compared with diffuse GFP fluorescence from the broadly labeled control. Matched zoomed regions at the SPress plane show resolved cellular structures in the SPress condition but not in the GFP control.

## References

1. K. D. Harris, A. Thiele, Cortical state and attention. Nat Rev Neurosci 12, 509–523 (2011).

2. L. Muller, F. Chavane, J. Reynolds, T. J. Sejnowski, Cortical travelling waves: mechanisms and computational principles. Nat Rev Neurosci 19, 255–268 (2018).

3. G. Buzsaki, A. Draguhn, Neuronal oscillations in cortical networks. Science 304, 1926–1929 (2004).

4. H. H. Yang, F. St-Pierre, Genetically Encoded Voltage Indicators: Opportunities and Challenges. J Neurosci 36, 9977–9989 (2016).

5. Y. Xu, P. Zou, A. E. Cohen, Voltage imaging with genetically encoded indicators. Curr Opin Chem Biol 39, 1–10 (2017).

6. T. Knopfel, C. Song, Optical voltage imaging in neurons: moving from technology development to practical tool. Nat Rev Neurosci 20, 719–727 (2019).

7. J. B. Pawley, Handbook Of Biological Confocal Microscopy. (2006).

8. W. Denk, J. H. Strickler, W. W. Webb, Two-photon laser scanning fluorescence microscopy. Science 248, 73–76 (1990).

9. J. Platisa et al., High-speed low-light in vivo two-photon voltage imaging of large neuronal populations. Nat Methods 20, 1095–1103 (2023).

10. T. D. Weber, M. V. Moya, K. Kilic, J. Mertz, M. N. Economo, High-speed multiplane confocal microscopy for voltage imaging in densely labeled neuronal populations. Nat Neurosci 26, 1642–1650 (2023).

11. L. Bai et al., Volumetric voltage imaging of neuronal populations in the mouse brain by confocal light-field microscopy. Nat Methods 21, 2160–2170 (2024).

12. S. Xiao et al., Large-scale deep tissue voltage imaging with targeted-illumination confocal microscopy. Nat Methods 21, 1094–1102 (2024).

13. J. Zhong et al., FACED 2.0 enables large-scale voltage and calcium imaging in vivo. Nat Methods, (2025).

14. Z. Wang et al., High-speed optical microscopy for neural voltage imaging: Methods, trade-offs, and opportunities. arXiv preprint arXiv:2512.15587, (2026).

15. S. Xiao et al., Large-scale voltage imaging in behaving mice using targeted illumination. iScience 24, 103263 (2021).

16. T. W. Chen et al., Imaging neuronal voltage beyond the scattering limit. Nat Methods 22, 1366–1375 (2025).

17. T. Saito, N. Nakatsuji, Efficient gene transfer into the embryonic mouse brain using in vivo electroporation. Dev Biol 240, 237–246 (2001).

18. L. Madisen et al., A robust and high-throughput Cre reporting and characterization system for the whole mouse brain. Nat Neurosci 13, 133–140 (2010).

19. L. T. Graybuck et al., Enhancer viruses for combinatorial cell-subclass-specific labeling. Neuron 109, 1449–1464 e1413 (2021).

20. S. Yao et al., RecV recombinase system for in vivo targeted optogenomic modifications of single cells or cell populations. Nat Methods 17, 422–429 (2020).

21. A. S. Abdelfattah et al., Sensitivity optimization of a rhodopsin-based fluorescent voltage indicator. Neuron 111, 1547–1563 e1549 (2023).

22. M. Steriade, A. Nunez, F. Amzica, A novel slow (< 1 Hz) oscillation of neocortical neurons in vivo: depolarizing and hyperpolarizing components. J Neurosci 13, 3252–3265 (1993).

23. S. Crochet, C. C. Petersen, Correlating whisker behavior with membrane potential in barrel cortex of awake mice. Nat Neurosci 9, 608–610 (2006).

24. J. F. Poulet, C. C. Petersen, Internal brain state regulates membrane potential synchrony in barrel cortex of behaving mice. Nature 454, 881–885 (2008).

25. I. Lampl, I. Reichova, D. Ferster, Synchronous membrane potential fluctuations in neurons of the cat visual cortex. Neuron 22, 361–374 (1999).

26. J. N. MacLean, B. O. Watson, G. B. Aaron, R. Yuste, Internal dynamics determine the cortical response to thalamic stimulation. Neuron 48, 811–823 (2005).

27. A. Luczak, P. Bartho, S. L. Marguet, G. Buzsaki, K. D. Harris, Sequential structure of neocortical spontaneous activity in vivo. Proc Natl Acad Sci U S A 104, 347–352 (2007).

28. M. H. Mohajerani et al., Spontaneous cortical activity alternates between motifs defined by regional axonal projections. Nat Neurosci 16, 1426–1435 (2013).

29. Y. Liang et al., Cortex-Wide Dynamics of Intrinsic Electrical Activities: Propagating Waves and Their Interactions. J Neurosci 41, 3665–3678 (2021).

30. W. E. Skaggs, B. L. McNaughton, M. A. Wilson, C. A. Barnes, Theta phase precession in hippocampal neuronal populations and the compression of temporal sequences. Hippocampus 6, 149–172 (1996).

31. D. J. Foster, M. A. Wilson, Reverse replay of behavioural sequences in hippocampal place cells during the awake state. Nature 440, 680–683 (2006).

32. K. D. Harris, G. M. Shepherd, The neocortical circuit: themes and variations. Nat Neurosci 18, 170–181 (2015).

33. H. Adesnik, A. Naka, Cracking the Function of Layers in the Sensory Cortex. Neuron 100, 1028–1043 (2018).

34. W. J. Zhao, J. Kremkow, J. F. Poulet, Translaminar Cortical Membrane Potential Synchrony in Behaving Mice. Cell Rep 15, 2387–2399 (2016).

35. M. L. Andermann et al., Chronic cellular imaging of entire cortical columns in awake mice using microprisms. Neuron 80, 900–913 (2013).

36. R. P. Barretto et al., Time-lapse imaging of disease progression in deep brain areas using fluorescence microendoscopy. Nat Med 17, 223–228 (2011).

37. Y. Senzai, A. Fernandez-Ruiz, G. Buzsaki, Layer-Specific Physiological Features and Interlaminar Interactions in the Primary Visual Cortex of the Mouse. Neuron 101, 500–513 e505 (2019).

38. M. V. Sanchez-Vives, D. A. McCormick, Cellular and network mechanisms of rhythmic recurrent activity in neocortex. Nat Neurosci 3, 1027–1034 (2000).

39. S. Sakata, K. D. Harris, Laminar structure of spontaneous and sensory-evoked population activity in auditory cortex. Neuron 64, 404–418 (2009).

40. N. J. Sofroniew, D. Flickinger, J. King, K. Svoboda, A large field of view two-photon mesoscope with subcellular resolution for in vivo imaging. Elife 5, (2016).

41. T. W. Chen et al., Ultrasensitive fluorescent proteins for imaging neuronal activity. Nature 499, 295–300 (2013).

42. J. S. Marvin et al., An optimized fluorescent probe for visualizing glutamate neurotransmission. Nat Methods 10, 162–170 (2013).

43. T. Patriarchi et al., Ultrafast neuronal imaging of dopamine dynamics with designed genetically encoded sensors. Science 360, (2018).

44. F. Sun et al., A Genetically Encoded Fluorescent Sensor Enables Rapid and Specific Detection of Dopamine in Flies, Fish, and Mice. Cell 174, 481–496 e419 (2018).

45. C. A. Saladrigas et al., Miniaturized widefield microscope for high speed in vivo voltage imaging. Biomed Opt Express 17, 1–11 (2026).

46. M. Kannan et al., Dual-polarity voltage imaging of the concurrent dynamics of multiple neuron types. Science 378, eabm8797 (2022).

47. H. Tian et al., Video-based pooled screening yields improved far-red genetically encoded voltage indicators. Nat Methods, (2023).

48. S. Inagaki et al., Isotonic and minimally invasive optical clearing media for live cell imaging ex vivo and in vivo. Nat Methods 23, 839–853 (2026).

49. M. E. Xie et al., High-fidelity estimates of spikes and subthreshold waveforms from 1-photon voltage imaging in vivo. Cell Rep 35, 108954 (2021).

50. K. Morikawa et al., Photoactivatable Cre recombinase 3.0 for in vivo mouse applications. Nat Commun 11, 2141 (2020).

51. J. B. Grimm et al., Bright photoactivatable fluorophores for single-molecule imaging. Nat Methods 13, 985–988 (2016).

52. M. P. Chien et al., Photoactivated voltage imaging in tissue with an archaerhodopsin-derived reporter. Sci Adv 7, (2021).

53. L. E. Fenno et al., Targeting cells with single vectors using multiple-feature Boolean logic. Nat Methods 11, 763–772 (2014).

54. J. Wu et al., A non-invasive far-red light-induced split-Cre recombinase system for controllable genome engineering in mice. Nat Commun 11, 3708 (2020).

55. A. S. Abdelfattah et al., Bright and photostable chemigenetic indicators for extended in vivo voltage imaging. Science 365, 699–704 (2019).

